# Assessment of Gene Set Enrichment Analysis using curated RNA-seq-based benchmarks

**DOI:** 10.1101/2024.01.10.575094

**Authors:** Julián Candia, Luigi Ferrucci

## Abstract

Pathway enrichment analysis is a ubiquitous computational biology method to interpret a list of genes (typically derived from the association of large-scale omics data with phenotypes of interest) in terms of higher-level, predefined gene sets that share biological function, chromosomal location, or other common features. Among many tools developed so far, Gene Set Enrichment Analysis (GSEA) stands out as one of the pioneering and most widely used methods. Although originally developed for microarray data, GSEA is nowadays extensively utilized for RNA-seq data analysis. Here, we quantitatively assessed the performance of a variety of GSEA modalities and provide guidance in the practical use of GSEA in RNA-seq experiments. We leveraged harmonized RNA-seq datasets available from The Cancer Genome Atlas (TCGA) in combination with large, curated pathway collections from the Molecular Signatures Database to obtain cancer-type-specific target pathway lists across multiple cancer types. We carried out a detailed analysis of GSEA performance using both gene-set and phenotype permutations combined with four different choices for the Kolmogorov-Smirnov enrichment statistic. Based on our benchmarks, we conclude that the classic/unweighted gene-set permutation approach offered comparable or better sensitivity-vs-specificity tradeoffs across cancer types compared with other, more complex and computationally intensive permutation methods. Finally, we analyzed other large cohorts for thyroid cancer and hepatocellular carcinoma. We utilized a new consensus metric, the Enrichment Evidence Score (EES), which showed a remarkable agreement between pathways identified in TCGA and those from other sources, despite differences in cancer etiology. This finding suggests an EES-based strategy to identify a core set of pathways that may be complemented by an expanded set of pathways for downstream exploratory analysis. This work fills the existing gap in current guidelines and benchmarks for the use of GSEA with RNA-seq data and provides a framework to enable detailed benchmarking of other RNA-seq-based pathway analysis tools.

## 1 Introduction

Since its inception two decades ago, pathway enrichment analysis has grown in popularity to become an essential data analysis tool in omics-based studies. By casting gene-level measurements in a broader biological context, this approach allows researchers to interpret their data in terms of a great variety of gene sets that may represent different functions, processes, components, or associations with disease. The success of pathway enrichment analysis, however, led to a very complex state of affairs with more than 70 different methods and hundreds of variants published to date [1–4] compounded by more than 33,000 human gene sets available from the Molecular Signatures Database (MSigDB) [5] alone, a scenario that has become extremely difficult to navigate.

Although pathway enrichment analysis methods share a common goal, their approach and underlying statistical models vary substantially. Existing methods have been categorized in various ways; for the purpose of our work, we are primarily concerned with over-representation analysis (ORA), whose input is an unranked list of genes (selected from the full list of measured genes by imposing a threshold criterion), and functional class scoring (FCS), whose input is the full list of measured genes (ranked based on quantitative scores) [6]. In recent years, several groups have discussed and implemented benchmarks to compare findings from different tools as well as to evaluate multiple factors that affect the results of enrichment analysis [1, 7–15]. Typically, these benchmarking efforts were aimed at comparing many different tools belonging to one or more broad categories, such as ORA and FCS. In contrast, our aim is to focus on the pioneering FCS-type method known as Gene Set Enrichment Analysis (GSEA) [16, 17], which arguably remains the most highly cited and widely used pathway analysis method, thereby comprehensively and thoroughly exploring its performance under different modalities by means of high-quality, curated RNA-seq datasets and pathway collections. Similarly to other early-generation pathway analysis methods, GSEA was originally developed for use with microarray data, but nowadays it is extensively used for analysis of RNA-seq data. Different omics technologies, however, are known to introduce a variety of potential sources of bias that may confound functional enrichment analysis [18, 19]. Thus, here we aim to assess GSEA’s performance specifically using the type of data on which GSEA is most commonly being currently utilized.

To that end, we leveraged an approach first proposed by Tarca et al. [7] and followed by several studies [1, 8, 12, 14, 15], which built a set of positive-control pathways using annotations that matched the conditions under study. In our approach, we scanned 11,274 harmonized RNA-seq samples across 33 cancer types from The Cancer Genome Atlas (TCGA) [20]. We applied filters to remove technical sources of variability and, in order to control for intertumor transcriptional heterogeneity, we focused on TCGA projects with available paired primary-tumor / non-tumor tissue samples. The final analysis involved 1,219 paired samples from 604 subjects across 12 cancer types. To select positive-control pathways, we scanned 33,591 gene sets from MSigDB (which includes KEGG [21], Reactome [22], Gene Ontology [23], and many other public resources) and implemented a sequence of filters leading to a total of 253 pathways associated to those 12 cancer types. GSEA was run using the latest available version 4.3.2 (build 13, October 2022) [24]. We generated random gene set ensembles to quantitatively assess the sensitivity and specificity of different GSEA modalities. Finally, we analyzed other large cohorts (a thyroid cancer study consisting of 780 paired tumor/nontumor RNA-seq tissue samples from 390 subjects [25] and a hepatocellular carcinoma study consisting of 140 paired tumor/nontumor RNA-seq tissue samples from 70 subjects [26]). By comparing results from these cohorts and their TCGA counterparts, we propose a new consensus metric termed Enrichment Evidence Score (EES) that allows the identification of a core set of pathways (those showing a high degree of consensus across enrichment metrics) complemented by an expanded set of pathways for exploratory analysis. Moreover, we show how the EES metric may be utilized to identify high-consensus leading-edge genes and integrate pathway- and gene-level information.

In summary, our primary goal is to leverage large, harmonized RNA-seq data repositories and large, curated pathway repositories in order to quantitatively assess the performance of a variety of GSEA modalities and provide guidance in the practical use of GSEA for the analysis of RNA-seq data. Moreover, a secondary goal of this study is to set up a framework capable of benchmarking other RNA-seq-based pathway analysis tools. To enable such secondary aim, we made all source code openly and publicly available along with detailed step-by-step documentation [27].

## 2 Methods

### 2.1 RNA-seq datasets

#### 2.1.1. The Cancer Genome Atlas (TCGA)

From NCI’s Genomic Data Commons portal [28] we downloaded all of TCGA’s gene expression quantification files and associated metadata. In total, these are 11,274 harmonized RNA-seq expression data files across 33 cancer types. In order to control for intertumor transcriptional heterogeneity, we only considered individuals with paired primary-tumor/non-tumor solid tissue samples available. Moreover, to remove technical sources of variability [29, 30], we excluded Formalin-Fixed Paraffin-Embedded (FFPE) tissue samples and considered only TCGA studies with at least 10 subjects available after these exclusions. This process left us with 15 TCGA projects. However, 3 of them (KICH, KIRP and UCEC) were later excluded because we could not identify enough positive-control pathways based on the procedure explained below (see Sect 2.3.1). The final analysis involved 1,219 paired primary-tumor/non-tumor solid tissue samples from 604 subjects across 12 cancer types (Table 1).

**Table 1.** Summary of TCGA projects and associated cancer-type-specific pathways from MSigDB.

| TCGA Project |  | RNA-seq Data |  | MSigDB Pathways |  |  |
| --- | --- | --- | --- | --- | --- | --- |
| ID | Description | Samples | Subjects | Preselected | Target | Positive-control |
| BLCA | Bladder urothelial carcinoma | 39 | 19 | 16 | 15 | 12 |
| BRCA | Breast invasive carcinoma | 228 | 113 | 137 | 109 | 72 |
| COAD | Colon adenocarcinoma | 84 | 41 | 30 | 25 | 15 |
| ESCA | Esophageal carcinoma | 26 | 13 | 10 | 7 | 5 |
| HNSC | Head and neck squamous cell carcinoma | 86 | 43 | 9 | 9 | 8 |
| KIRC | Kidney renal clear cell carcinoma | 144 | 72 | 8 | 6 | 3 |
| LIHC | Liver hepatocellular carcinoma | 100 | 50 | 114 | 92 | 70 |
| LUAD | Lung adenocarcinoma | 122 | 58 | 39 | 29 | 16 |
| LUSC | Lung squamous cell carcinoma | 102 | 51 | 39 | 29 | 16 |
| PRAD | Prostate adenocarcinoma | 104 | 52 | 27 | 24 | 16 |
| STAD | Stomach adenocarcinoma | 66 | 33 | 12 | 11 | 8 |
| THCA | Thyroid carcinoma | 118 | 59 | 30 | 21 | 12 |

#### 2.1.2 Thyroid cancer from the Chernobyl Tissue Bank (REBC-THYR)

Following a similar procedure as described above, we used NCI’s Genomic Data Commons portal [28] to download gene expression quantification files and associated metadata for REBC-THYR, a radiation-induced thyroid cancer study from the Chernobyl Tissue Bank [25]. Based on the same exclusion criteria followed for TCGA data, we obtained a total of 780 paired primary-tumor/non-tumor solid tissue samples from 390 subjects.

#### 2.1.3 Hepatocellular carcinoma from Mongolia (MO-HCC)

From NCBI’s Gene Expression Omnibus repository (accession GSE144269) [31], we downloaded gene expression quantification files and associated metadata from a study of hepatocellular carcinoma in Mongolia, which comprises a total of 140 paired primary-tumor/non-tumor solid tissue samples obtained from 70 subjects [26].

### 2.2 Differential Gene Expression (DGE) analysis

All of TCGA’s gene expression quantification files are openly available to download from NCI’s Genomic Data Commons (GDC) portal [28]; we provide step-by-step details in our code and documentation repository [27]. The GDC mRNA quantification analysis pipeline measures gene level expression with STAR as raw read counts.

Subsequently, the counts are augmented with several transformations including Fragments per Kilobase of transcript per Million mapped reads (FPKM), upper quartile normalized FPKM (FPKM-UQ), and Transcripts per Million (TPM). These values are additionally annotated with the gene symbol and gene bio-type. These data are generated through this pipeline by first aligning reads to the GRCh38 reference genome and then by quantifying the mapped reads. To facilitate harmonization across samples, all RNA-Seq reads are treated as unstranded during analyses. Further details are provided in the GDC documentation [32].

Gene expression count matrices consist of 60,660 quantified genes, out of which we kept 19,962 protein-coding genes. After correcting for gene name aliases, we found that 19,321 of these genes were included in pathways from MSigDB. We then performed further filtering and DGE analysis separately for each TCGA project. Using TPM quantification, we removed low-abundance genes with TPM *>* 1 in less than 40% of the samples, transformed the count matrix using log_2_ (TPM + 1) and fitted the expression values against tissue type (primary tumor or non-tumor tissue) and subject (to take into account the paired sample design) using functions lmFit, eBayes and topTable from the limma [33] R package v.3.56.1. S1 Table shows the number of measured genes in each TCGA project after all filters were applied and how many of them were found differentially expressed in either direction, i.e. over-expressed in primary-tumor (fold change *FC >* 1) or non-tumor (*FC <* 1) tissue, using Bonferroni or Benjamini-Hochberg (B-H) − q value < 0.05 significance thresholds adjusted for multiple testing. An illustration of these results is shown in S1 Fig with a Volcano plot of significant genes based on Bonferroni-adjusted q − value < 0.05 for TCGA-BRCA (breast cancer).

In order to explore an alternative procedure for count normalization and gene filtering feeding into the DGE analysis, we re-analyzed all paired primary-tumor vs non-tumor cancer samples from TCGA using raw counts. We implemented a standard analysis pipeline using the edgeR package v.3.42.2 by applying filterByExpr to filter genes, followed by voom quantile normalization and limma for assessing differentially expressed genes with variance stabilization. Using standard gene ranking scores, defined by Eq.( 3) below, S2 Fig illustrates the excellent agreement between the two procedures for TCGA-BRCA samples. Furthermore, in order to extend the comparison to the pathway level, we re-ran GSEA with gene set permutations (described below, see Sect. 2.4 for details) using the voom-normalized data and determined pathway-level ranking scores based on GSEA p-values and enrichment directions. S3 Fig shows comparisons of these pathway-level ranking scores derived from the two alternative count normalization procedures for unweighted (classic) and weighted (*p* = 1, 1.5 and 2) enrichment statistics across all TCGA-BRCA target pathways (see below for further details). Taken together, S2 Fig and S3 Fig show that differential expression/enrichment analyses derived from these different count normalization and filtering procedures lead to highly concordant results at both gene and pathway levels.

### 2.3 Gene sets

#### 2.3.1 Cancer-type-specific pathways

Fig 1 illustrates the process to obtain cancer-type-specific pathways as positive controls. We downloaded 33,591 human gene sets from MSigDB v2023.1.Hs (released on March 2023) [34]. Using functionality provided by the R package fgsea v.1.26.20, we ran a custom script to identify, for each cancer type, gene sets whose names matched a curated list of search terms. The preselected pathways were then ran through a size filter procedure (following GSEA recommendations, we excluded gene sets outside the 15-500 size range, considering only genes that passed all filters in the quantification step). After running GSEA in multiple modalities, we applied a subsequent filter to keep target pathways that were found significant (unadjusted p − value < 0.05) at least once after running GSEA with gene-set and phenotype permutations using all available enrichment statistic settings. Table 1 shows the number of pathways selected at each stage. Full details are provided in S2 Table.

**Fig 1.**
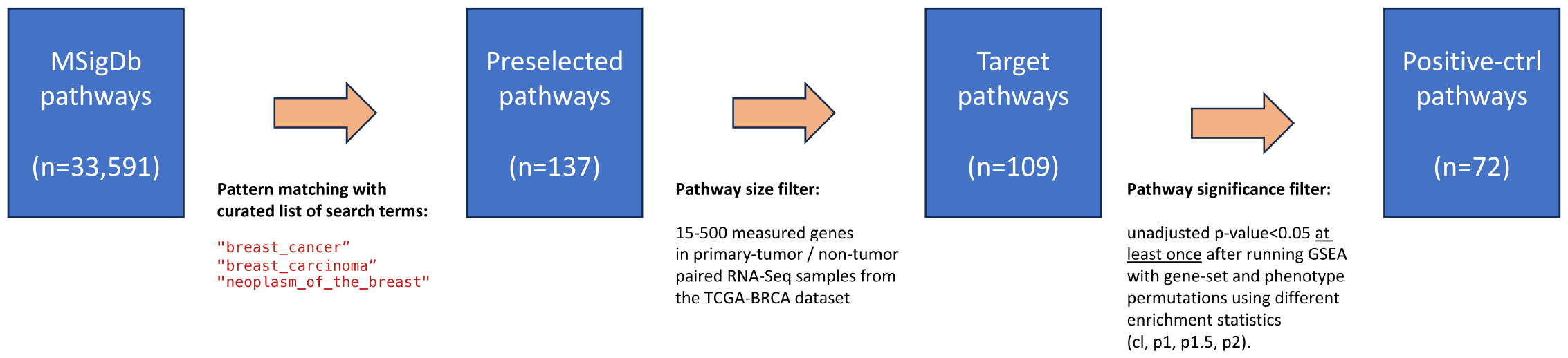
Cancer-type-specific pathways. Procedure to identify lists of preselected, target, and positive-control pathways using TCGA-BRCA (breast cancer) as example. See S2 Table for full details.

#### 2.3.2 Gene set overlap

An important challenge of pathway enrichment analysis is that of gene set overlap, where some genes participate in multiple gene sets [35, 36]. This phenomenon occurs in the presence of multifunctional genes (i.e., genes that play a role in several biological functions or molecular processes) but is also prevalent in some gene set collections with redundancies or a strong hierarchical structure. The latter case is most notably present, by design, in the three lineages of Gene Ontology (GO) [23]. A useful metric used to quantify similarities between gene sets 𝒢_*i*_ and 𝒢_*j*_ is the Jaccard index, given by

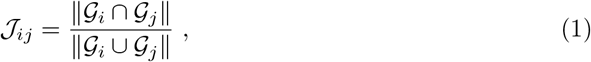

which defines a symmetric matrix normalized to the 0 − 1 range. Alternatively, the fractional pairwise overlap matrix, given by

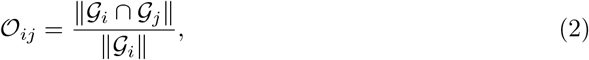

defines a non-symmetric matrix normalized to the 0 − 1 range that better captures hierarchical relations such as those present in GO.

### 2.4 Gene Set Enrichment Analysis (GSEA)

Gene Set Enrichment Analysis is a well-established tool based on a weighted, normalized Kolmogorov-Smirnov statistic to assess pathway enrichment of ranked gene lists [16, 17]. Originally developed for microarray data, GSEA is a threshold-free method that analyzes all genes on the basis of their ranking score, without prior gene filtering, and is thus recommended when ranks are available for all or most of the genes in the genome (e.g., transcriptomics) [37].

Let *r*_*k*_ be the ranking score associated with the *k* − th gene, *g*_*k*_, with *k* = 1, …, *n* an index running through all measured genes. Without loss of generality, we assume genes to be indexed in decreasing order of their ranking scores. Typically, it is assumed that ranking scores span positive and negative values; the absolute value of a ranking score is a measure of effect size and/or statistical significance, whereas the sign indicates the direction of change. For a case-control differential gene expression design with fold change *FC*, one popular choice is a ranking score of the form

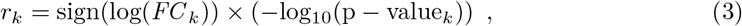

where p − value_*k*_ is an estimate of the statistical significance of the observed *FC*_*k*_ for the *k* − th gene. Based on the DGE analysis outlined above (Sect. 2.2), we obtained standard ranking scores, defined by Eq.( 3), which are provided in S3 Table for all TCGA projects analyzed in our study.

Let 𝒢_*i*_ be a gene set (after removing any genes not included in the universe of measured genes). The step variable *x*_*k*_ is defined as

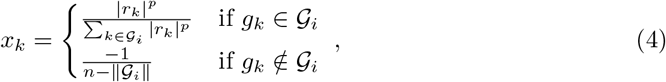

where *p >* 0 is a weight parameter introduced in GSEA’s follow-up paper [17] to emphasize the role of genes at the top and bottom of the gene list (i.e. those with larger absolute values of the ranking score in either direction) in detriment of genes in the center of the list (i.e. those with lower absolute values of the ranking score in either direction). The denominators are chosen for normalization purposes. The *p* = 0 case is akin to the standard, unweighted formulation from GSEA’s original paper [16], although with a different normalization.

The cumulative enrichment of gene set 𝒢_*i*_ is defined as the running sum of the step variable *x*_*k*_ from top to bottom, i.e.

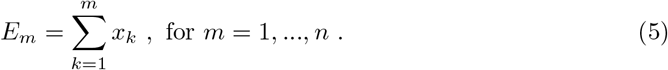

In this way, starting at *E*_0_ ≡ 0, GSEA progressively examines genes from the top to the bottom of the ranked list, increasing the cumulative enrichment if the gene at that position is part of the pathway and decreasing the running sum otherwise. The step variable is normalized to ensure that *E*_*n*_ = 0 at the bottom of the list. Then, the target gene set’s enrichment score (*ES*) is defined as the maximum departure from zero (in either direction, i.e. positive or negative) along the cumulative enrichment profile; its sign indicates whether the observed enrichment is associated with the top (positive *ES*) or bottom (negative *ES*) of the ranked gene list. Genes in the target gene set that contribute to the maximum departure from zero in the cumulative enrichment running sum are termed leading-edge or core-enrichment genes. By definition of the step variable normalization, *ES* is restricted to the [ − 1, 1] range; the extreme *ES* = ± 1 values may only occur when all genes in a gene set appear strictly at the top (*ES* = 1) or bottom (*ES* = −1) of the ranked gene list. Because, by virtue of Eqs.(4)-(5), GSEA’s weighted Kolmogorov-Smirnov (global) statistic cannot be expressed in terms of a simple function of the ranking score (local statistic), formal analytical solutions to express the significance of a pathway’s enrichment score based on parametric modeling are not known. Instead, non-parametric methods based on empirical distributions are computationally generated via random permutations. GSEA offers two options: (i) gene-set permutations, in which a target gene set’s enrichment score is compared against a null-model distribution of enrichment scores from an ensemble of randomly generated gene sets of equal size, and (ii) phenotype permutations, in which the null-model distribution is generated by randomly shuffling phenotype labels among the samples while keeping the original target gene set.

GSEA’s GUI applications offer a limited choice of ranking statistics and design matrices [37], which are not appropriate in the context of our work (see Sect. 4 below for a detailed discussion). For gene-set permutations, it is possible to upload the ranked list of genes into GSEA’s GUI in Preranked mode; however, GSEA does not offer suitable functionality to perform phenotype permutations on RNA-seq data [37].

Because our study involved both gene-set and phenotype permutations, we implemented GSEA in the command line via custom scripts. In this work, GSEA was run using `gsea-cli.sh` in a macOS environment (Monterey v.12.6.9) from the latest available GSEA version 4.3.2 (build 13, October 2022) [24]. Importantly, it must be pointed out that `gsea-3.0.jar`, utilized in protocols published by Reimand et al [37], is affected by serious security vulnerabilities due to the use of the Java-based logging utility Apache Log4j in GSEA versions earlier than 4.2.3. Moreover, as reported by the GSEA Team, version 3.0 contained microarray-specific code (mostly related to Affymetrix) that may cause issues with RNA-seq data analysis, which was removed in later GSEA updates.

Our analysis of phenotype permutations followed an approach similar to that outlined by Reimand et al. [37]. For each TCGA project under study, we generated *n*_*rdm*_ = 1000 phenotype randomizations and obtained *n*_*rdm*_ randomized list of ranks, which were distributed among parallel command-line GSEA executions. The statistical significance for each pathway was then empirically estimated as

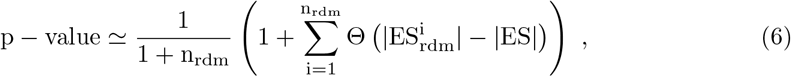

where Θ is the Heaviside step function. This expression computes the fraction of randomized ranked gene lists that yield enrichment scores 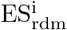 larger (in absolute value) than the enrichment score ES obtained in the original, non-randomized ranked gene list, correcting for random sampling [38]. It should be noticed that, because our matrix design was paired, the phenotype permutation procedure randomized tumor vs non-tumor labels while preserving subject identifiers [39].

In this work, we introduce a new consensus metric termed Enrichment Evidence Score (EES). At the pathway level, EES is defined as the signed sum of GSEA runs in which the pathway was significantly enriched, summed over the four enrichment statistics (classic and weighted with *p* = 1, 1.5, and 2). For instance, EES = 3 would correspond to a pathway found significantly enriched in tumor samples (positive sign) in three out of the four GSEA runs, whereas EES = − 2 would correspond to a pathway found significantly enriched in non-tumor samples (negative sign) in two out of the four GSEA runs. Similarly, we define a gene-level EES by summing over GSEA runs in which a given gene was identified as leading-edge.

### 2.5 Over-representation analysis (ORA)

ORA is the pioneer approach for pathway enrichment analysis, originally designed for microarray data [40]. Despite its simplicity, it is a robust and well-established method that has been implemented in more than 40 tools [41], which differ in their various components such as gene set database, data visualization, and user interface. Unlike GSEA, ORA requires a selection criterion to define a list of top ranking genes, ℒ, and treats all selected genes equally, ignoring measurable quantitative differences (e.g. the magnitude of gene expression differences). ORA tests the null hypothesis of no association between genes in ℒ and those contained in a target gene set 𝒢_*i*_. The probability for the observed intersection between these sets, *n*_*i*_ ≡ ∥ℒ ∩ 𝒢_*i*_ ∥, is given by the hypergeometric distribution, i.e.

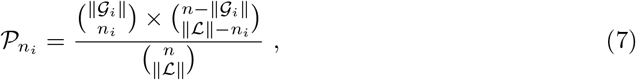

and the cumulative probability of an intersection equal or larger than that observed is given by

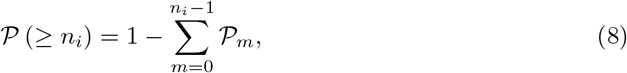

which matches the one-sided Fisher’s exact test of the association between genes in ℒ and 𝒢_*i*_. This approach lies at the core of many popular tools for pathway analysis, both public and proprietary, such as PathwayStudio [42], DAVID [43], g:Profiler [44] and Ingenuity Pathway Analysis [45], among many others [41].

We generated custom implementations of ORA using multiple modalities. After generating DGE tables with tumor/non-tumor *FC* and p − value estimates (see Sect. 2.2), we proceeded according to two different approaches, namely (i) signed ORA and (ii) unsigned ORA. In the signed ORA approach, we performed two analyses, one using significant genes with *FC* > 1 and another one using significant genes with *FC* < 1; a target pathway was thus assigned two enrichment p − values, which were merged by choosing the most significant between the two. In the unsigned ORA approach, in contrast, *FC* was ignored and genes were selected solely based on their p − values. Pan et al [46] pointed out one shortcoming of ORA, namely that pathways identified as statistically significant may strongly depend on the cutoffs used to select the input lists of differentially expressed genes. To take this effect into consideration, we implemented the q − value < 0.05 selection criterion applied to two multiple-testing correction scenarios, namely: (i) Bonferroni-adjusted p-values (stricter and therefore leading to shorter input gene lists), and (ii) Benjamini-Hochberg (also known as false discovery rate, FDR), as an alternative, more lenient approach used to select lists for ORA. In combination with the signed/unsigned ORA approaches mentioned above, these gene selection criteria yield a total of four ORA-based scenarios.

### 2.6 Analysis of sensitivity and specificity

To perform an analysis of sensitivity and specificity for each TCGA project in this study, we generated an ensemble of 1000 negative-control random pathways. Pathway sizes in the 15 − 500 range were sampled from a uniform distribution; genes were randomly chosen from the set of measured genes that passed the filters (see Sect. 2.2 for data processing details). After ordering positive- and negative-control pathways by enrichment p − value, we used the roc function from R package pROC [47] v.1.18.4 to generate one-directional receiver operating characteristic (ROC) curves to test the assumption that positive-control pathways are more significantly enriched than negative-control ones. The Area Under the Curve (AUC) was calculated with function auc and DeLong’s 95% confidence interval with function ci.auc from the same package. For visualizations, we used R package VennDiagram v.1.7.3, R package gplots v.3.1.3, and Cytoscape v.3.9.1.

## 3 Results

We begin to explore our ideas using TCGA-BRCA (breast cancer), the most common cancer type in the U.S. and worldwide, for which we have 228 paired primary-tumor/non-tumor RNA-seq samples available from 113 subjects. Following the procedure outlined in Sect. 2.3.1, a sequential series of filters was applied on MSigDB gene sets to identify 72 positive-control pathways (Fig 1). The analysis of gene set overlap metrics, namely the Jaccard index (Eq.(1), see S4 Fig) and the fractional pairwise overlap matrix (Eq.(2), see S5 Fig), revealed that positive-control pathways are mostly non-overlapping. We then ran GSEA under eight different modalities, choosing gene-set or phenotype permutations combined with four different choices for the enrichment statistic, either classic (unweighted) or weighted (*p* = 1, 1.5, 2), which comprise all the options available in the current GSEA release (see Sect. 2.4 for further details).

Fig 2(a) shows the overlap of positive-control pathways found significant among different weight parameter choices using gene-set permutations and p − value < 0.05 as significance criterion. We observe that, while half (36) of the pathways were found significant by all four runs and an additional set of 8 were found significant by three runs, 9 pathways were identified only by two methods and 19 of them were found by only one method. Among the latter, most of them (17) were identified by the classic, unweighted method. By tightening the significance criterion to p − value < 0.01, shown in Fig 2(b), the percentage of pathways found significant by all methods increases only slightly to 56% (32 out of 57 pathways). We find that stronger consensus cannot be enforced by tightening the significance criterion even further; using p − value < 0.001, still only 56% (26 out of 46) pathways agree among all methods (see S6 Fig). Similarly, Figs 2(c)-(d) show results for phenotype permutations using different p-value cutoffs, as indicated. It is clearly apparent that the phenotype permutation approach identifies fewer pathways and shows poorer agreement among different weight parameter choices. For stricter selection criteria (p − value < 0.001), none of the 72 positive control pathways are identified.

**Fig 2.**
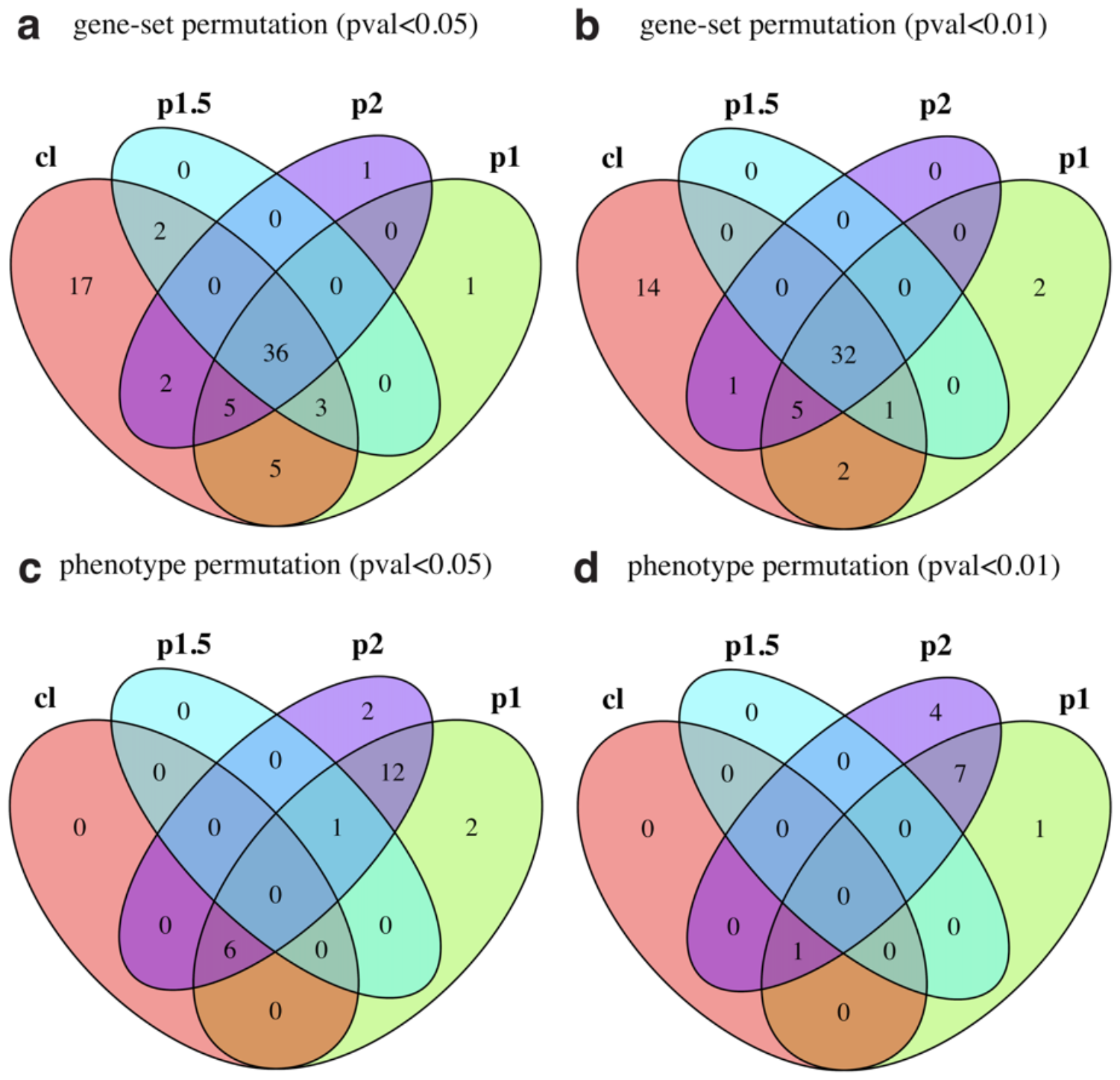
Significant TCGA-BRCA positive control pathways across different weight parameter choices. (a) Gene-set permutation with p − value < 0.05. (b) Gene-set permutation with p − value < 0.01. (c) Phenotype permutation with p − value < 0.05. (d) Phenotype permutation with p − value < 0.01. GSEA enrichment statistics: classic (“cl”), weight parameter *p* = 1 (“p1”), weight parameter *p* = 1.5 (“p1.5”), and weight parameter *p* = 2 (“p2”).

To complement these findings, Fig 3 shows the overlap between gene-set and phenotype permutation methods at significance p − value < 0.05 for each enrichment statistic. As shown in this figure, phenotype permutation is significantly less sensitive than gene-set permutation, since the former approach identifies subsets of less than half the number of positive control pathways obtained by the latter. Furthermore, to explore the impact of tumor/non-tumor sample pairedness, which affects the magnitude and statistical significance of differential gene expression estimates (and, therefore, gene ranking metrics) as well as phenotype randomization procedures (Sect. 2.4), we re-ran our pipeline as unpaired data. As shown by S7 Fig (for gene-set permutations) and S8 Fig (for phenotype permutations), differences between paired and unpaired analyses appear to be minor.

**Fig 3.**
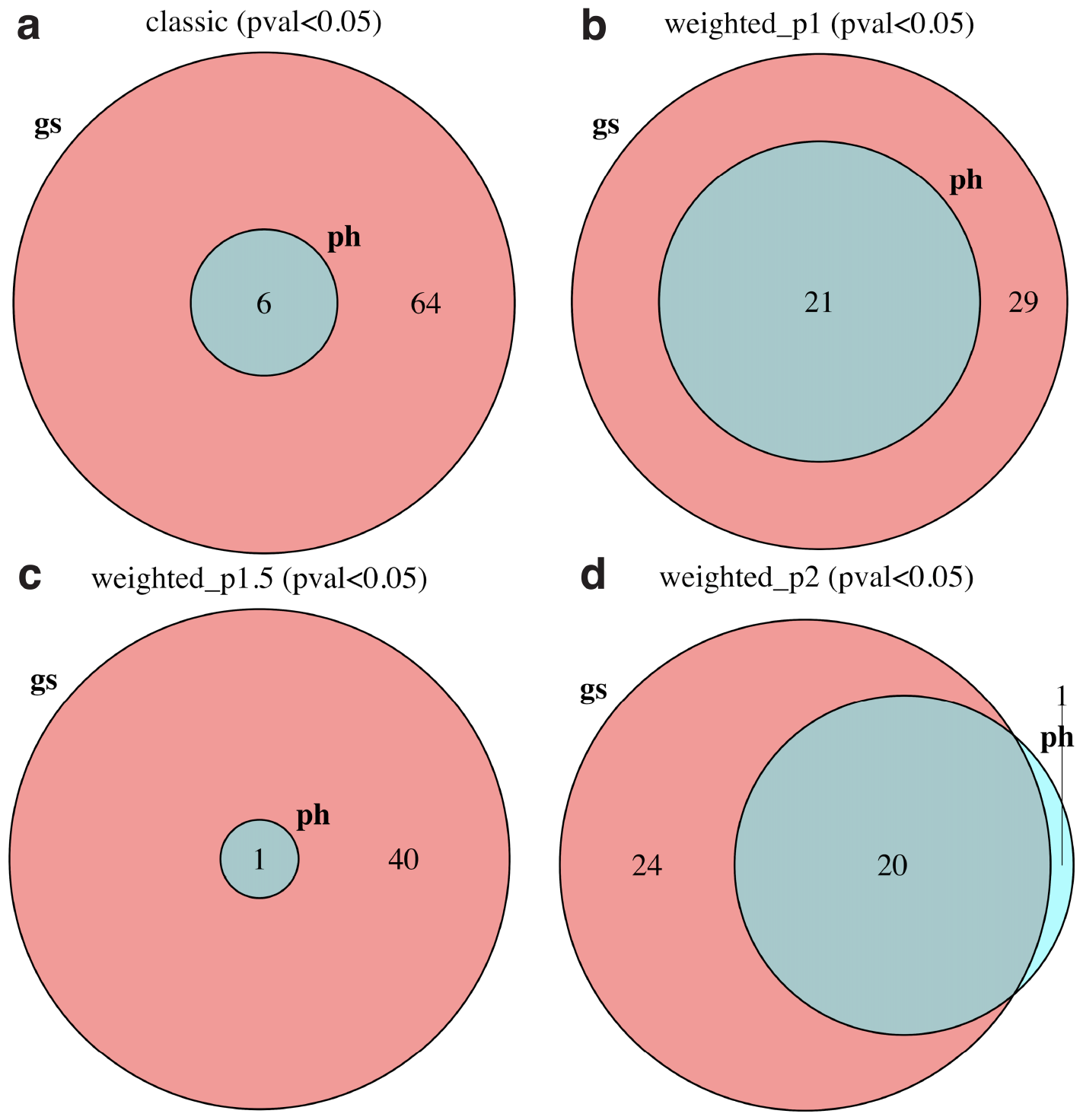
Significant TCGA-BRCA positive control pathways using gene-set (“gs”) and phenotype (“ph”) permutation approaches for different enrichment statistics. The significance criterion was p − value < 0.05. (a) Classic (unweighted). (b) Weight parameter *p* = 1. (c) Weight parameter *p* = 1.5. (d) Weight parameter *p* = 2.

In order to compare the sensitivity and specificity of GSEA under different modalities, we generated 1000 negative-control pathways by random selection of genes (see Sect. 2.6 for details). Ordering positive and negative controls by significance, Fig 4 shows receiver operating characteristic (ROC) curves for (a) gene-set and (b) phenotype permutation approaches, using different enrichment statistics. For comparison, we also show results from (c) signed and (d) unsigned implementations of over-representation analysis (ORA) using gene lists selected by q − value < 0.05 adjusted by two different methods (see Sect. 2.5 for details). This analysis shows that the classic gene-set permutation approach of GSEA yields the best sensitivity vs specificity tradeoff with Area Under the Curve (AUC) estimated as 0.99 and DeLong’s 95% confidence interval (CI) in the 0.98 − 1 range. The introduction of weighted enrichment statistics appears to be deleterious, negatively impacting the performance of gene-set approaches (Fig 4(a)). Moreover, in agreement with previous results, phenotype permutation models display poorer sensitivity (Fig 4(b)). The straightforward signed ORA (Fig 4(c)) achieves a performance similar to that of gene-set GSEA with *p* = 1 and better than that of other weighted approaches, but below gene-set classic GSEA. It should be noticed that our definition of positive-control pathways relies on a pathway significance filter, which removes target pathways that appeared not significant in any of the GSEA runs (Sect. 2.3.1 and Fig 1). By removing this filtering step and, thereby, expanding the target list from 72 to 109 pathways, our main conclusions remain the same: the classic gene-set permutation approach of GSEA is the best-performing model with AUC=0.87 (95% CI=0.83 − 0.91), as shown in S9 Fig.

**Fig 4.**
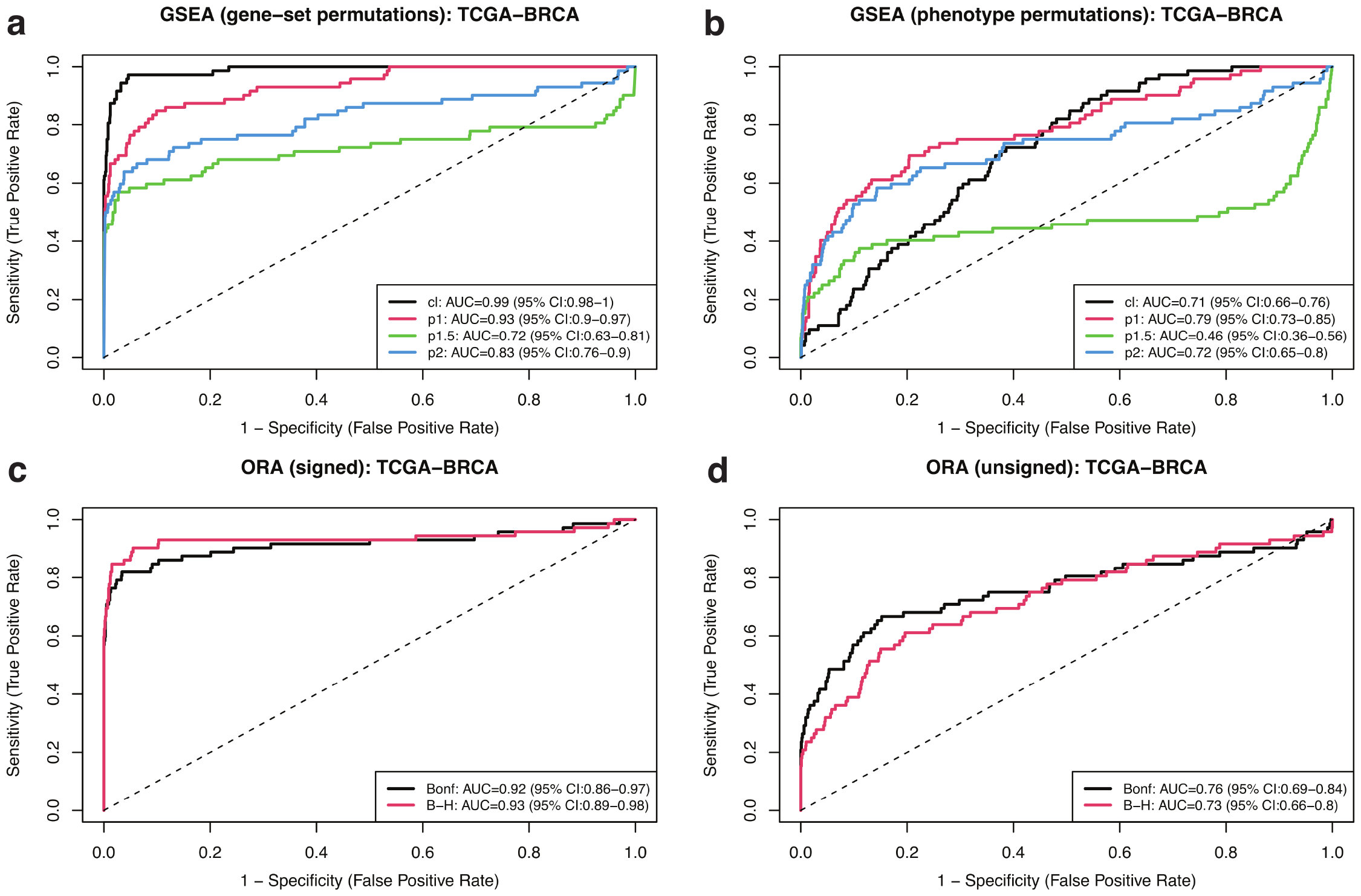
ROC curves for different GSEA and ORA approaches using 72 TCGA-BRCA positive control pathways and 1000 randomized negative controls. (a) Gene-set permutation GSEA. (b) Phenotype permutation GSEA. (c) Signed ORA. (d) Unsigned ORA. GSEA approaches used different enrichment statistics, as indicated. ORA approaches used Bonferroni and Benjamini-Hochberg (B-H) adjusted q-values as different inclusion criteria to select differentially expressed genes, as indicated.

Fig 5 expands the analysis by showing the Area Under the Curve across all cancer types included in our study, where error bars indicate DeLong’s 95% confidence intervals, for different GSEA and ORA approaches using positive control pathways and randomized negative controls as previously described. Despite the fact that the number of available samples, subjects, and pathways identified as positive controls varies significantly across projects (recall Table 1), the classic gene-set permutation approach of GSEA remains, quite remarkably, the top performing method. As shown in S10 Fig, this finding is confirmed by expanding the analysis to include all target pathways (i.e. by removing the last pathway filtering step in the procedure described in Sect. 2.3.1 and Fig 1).

**Fig 5.**
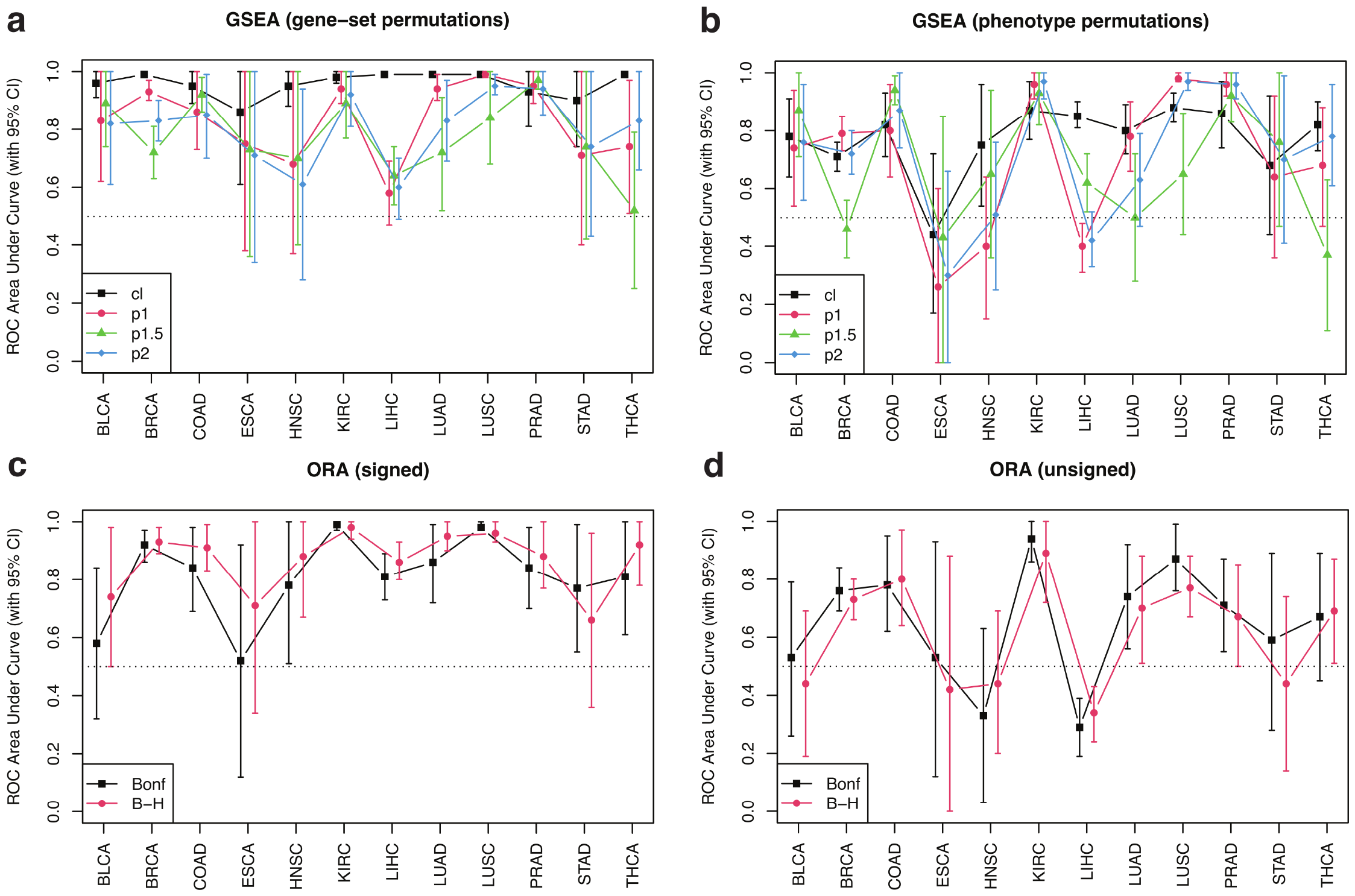
AUC across TCGA projects for different GSEA and ORA approaches using cancer-type-specific positive control pathways and 1000 randomized negative controls. (a) Gene-set permutation GSEA. (b) Phenotype permutation GSEA. (c) Signed ORA. (d) Unsigned ORA. GSEA approaches used different enrichment statistics, as indicated. ORA approaches used Bonferroni and Benjamini-Hochberg (B-H) adjusted q-values as different inclusion criteria to select differentially expressed genes, as indicated.

As discussed earlier (see Sect. 2.2), alternative methods for differential gene expression analysis exist, which rely on a variety of approaches for count normalization and statistical models for differential expression quantification. For the sake of comparison, S11 Fig shows ROC curves for GSEA gene-set permutations under different enrichment statistics obtained from TCGA-BRCA’s raw counts analyzed using the edgeR-voom-limma pipeline previously described. Although the differences among different enrichment statistics appear to be less pronounced compared with Fig 4(a), the unweighted Kolmogorov statistic approach still yields the largest AUC value. Similarly, S12 Fig extends the comparison to all TCGA projects analyzed in this study.

Finally, we explored similarities and differences between target pathways identified in TCGA versus other cancer-type-specific cohorts. Firstly, we analyzed REBC-THYR, a thyroid cancer study consisting of 780 paired tumor/nontumor RNA-seq tissue samples from 390 subjects (Sect. 2.1.2). Using gene-set permutations in GSEA with threshold p − value < 0.05, we calculated the Enrichment Evidence Score (EES) for each target pathway (see Sect. 2.4 for details). Fig 6(a) shows the contingency table of EES across target pathways for REBC-THYR vs TCGA-THCA, which displays a remarkable agreement between pathways detected in both thyroid cancer studies (see S4 Table for details). By grouping EES intervals as: (i) [ − 4, − 3] (non-tumor-enriched), (ii) [ − 2, 2] (undetermined), and (iii) [3, 4] (tumor-enriched), the association between enriched pathways is further emphasized, as shown in Fig 6(b), leading to Fisher’s exact test p − value = 2.5 × 10^−6^. Secondly, we analyzed MO-HCC, a study on hepatocellular carcinoma (the most common form of liver cancer) consisting of 140 paired tumor/nontumor RNA-seq tissue samples from 70 subjects (Sect. 2.1.3). Following a similar procedure, we obtained the contingency table of EES across target pathways for MO-HCC vs TCGA-LIHC, shown in Fig 6(c) and S5 Table, as well as the contingency table by grouped intervals, displayed in Fig 6(d). These results appear to capture the larger heterogeneity of TCGA-LIHC compared with MO-HCC, which is to be expected since the Mongolia study is focused on a cohort of rather uniform genetic background from a relatively isolated region exposed to weaker external influences [26]. Despite the spread attributable to differences in etiology, the association between EES groups appears highly significant, with Fisher’s exact test p − value = 4.4 × 10^−15^.

**Fig 6.**
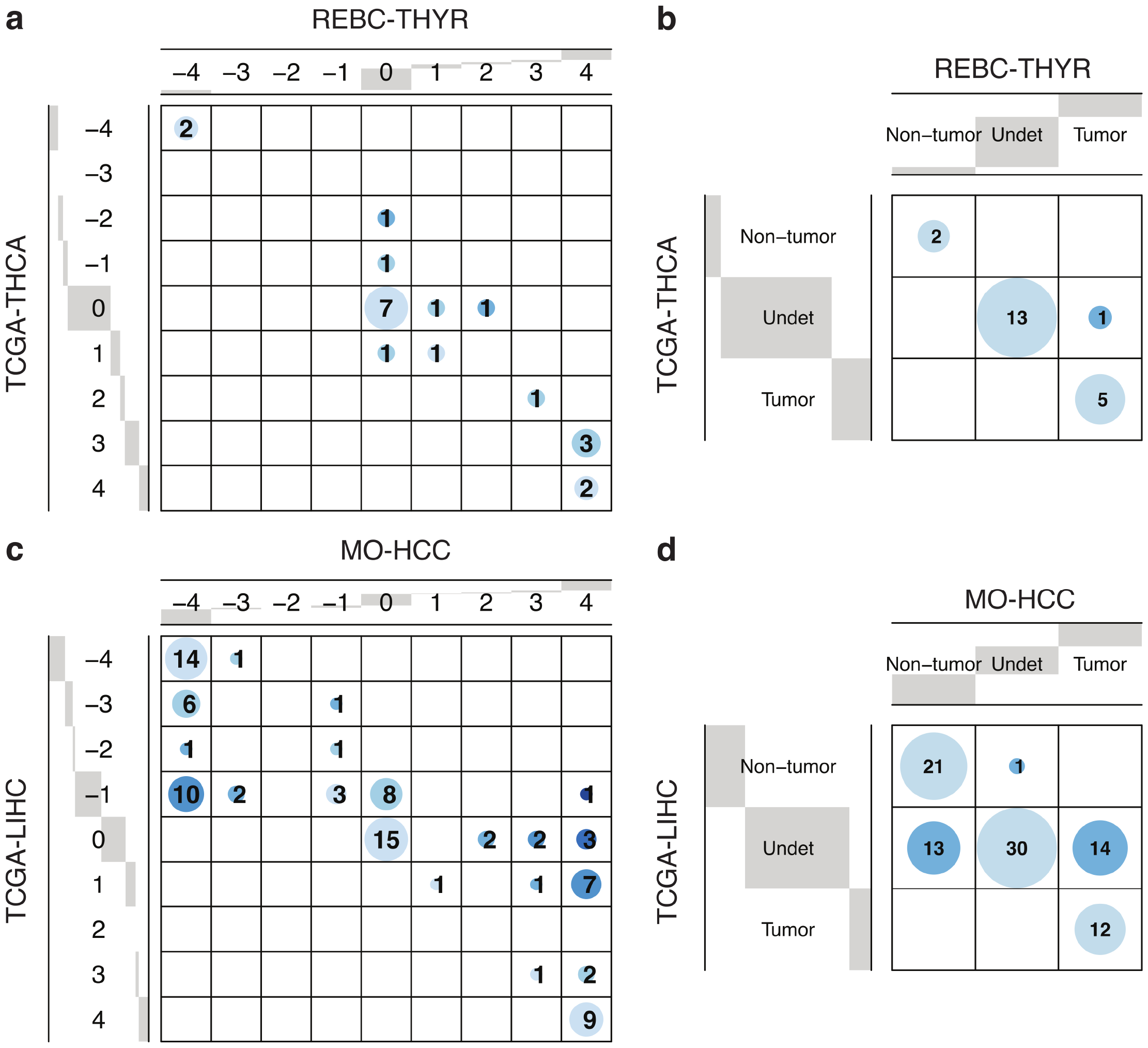
Pathway-level Enrichment Evidence Scores for thyroid cancer and hepatocellular carcinoma cohorts. (a) Comparison between significant target pathways in REBC-THYR vs TCGA-THCA. (b) Contingency table of grouped EES intervals in REBC-THYR vs TCGA-THCA (Fisher’s exact test p − value = 2.5 × 10^−6^). (c) Comparison between significant target pathways in MO-HCC vs TCGA-LIHC. (d) Contingency table of grouped EES intervals in MO-HCC vs TCGA-LIHC (Fisher’s exact test p − value = 4.4 × 10^−15^). Distance to the diagonal is represented with increasingly darker shades of blue.

A similar approach can also be implemented to investigate the role of leading-edge genes within the previously identified core pathways. Leading-edge genes are those identified by the GSEA algorithm to make the strongest contribution to the enrichment signal of a pathway (see Sect. 2.4 for details). For the sake of illustration, Fig 7(a) shows gene-level EES scores for LUI THYROID CANCER CLUSTER 1, a previously identified high-consensus tumor-enriched pathway in thyroid cancer (recall S4 Table).

**Fig 7.**
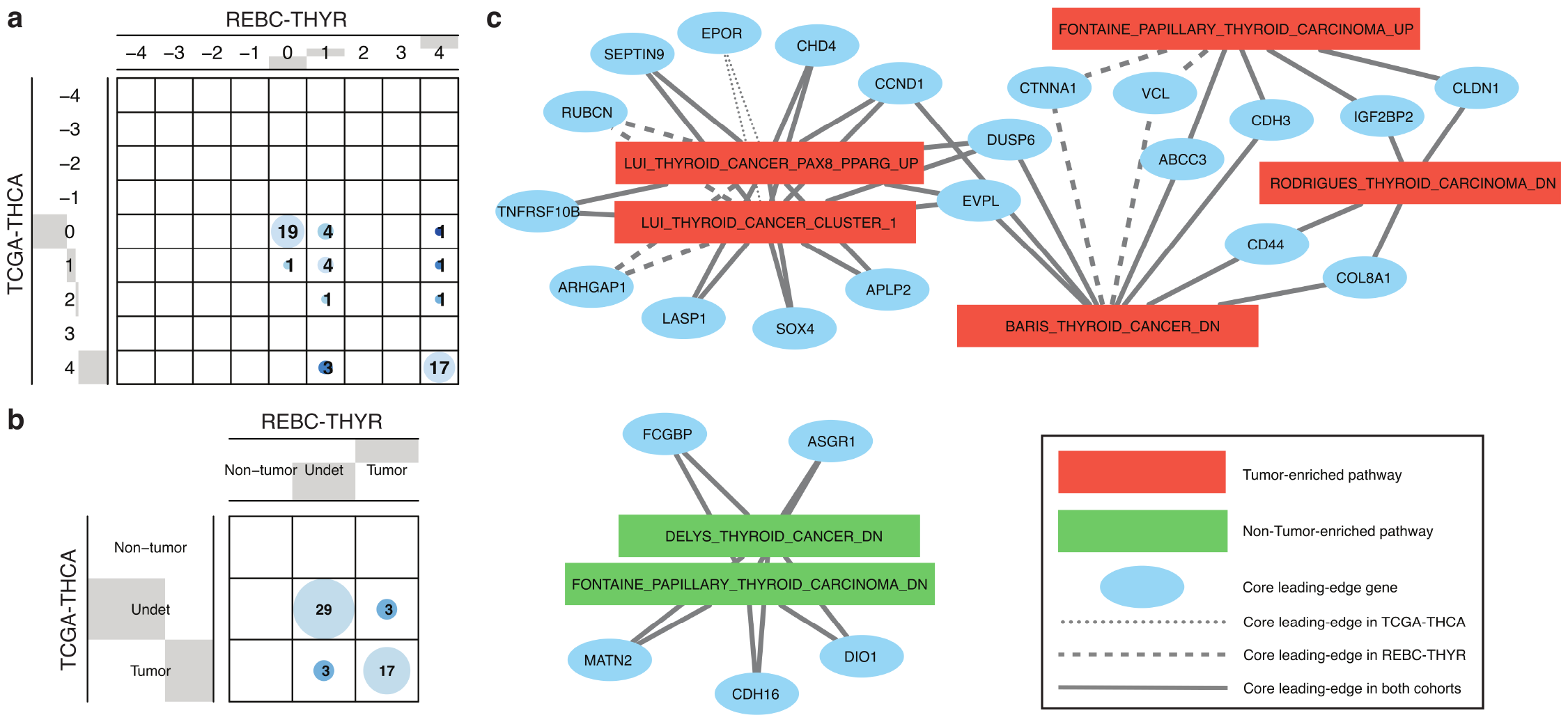
Gene-level Enrichment Evidence Scores for thyroid cancer cohorts. (a) Comparison between EES leading-edge genes in REBC-THYR vs TCGA-THCA for genes in LUI THYROID CANCER CLUSTER 1, previously identified as a high-consensus tumor-enriched pathway in thyroid cancer. (b) Contingency table of grouped EES intervals for the same case as in panel (a) (Fisher’s exact test p − value = 4.6 × 10^−8^). (c) Network representation showing seven core thyroid cancer pathways and high-consensus leading-edge genes ( |EES| ≥3 in at least one of the cohorts). Only genes connected to two or more pathways are shown.

The association between EES groups, shown in Fig 7(b), is highly significant, with Fisher’s exact test p − value = 4.6 × 10^−8^. Out of 52 constituent genes, we observe that 17 of them are robustly identified as leading-edge genes in this thyroid cancer pathway in both cohorts (see S6 Table for details). Interestingly, three genes (EPOR, GAS6, CDC25B) are found to be leading-edge only in TCGA-THCA, while other three genes (SPRY1, ARHGAP1, RUBCN) are identified as leading-edge only in REBC-THYR, which may point to mechanistic differences across etiologies. Fig 7(c) illustrates how we can integrate high-consensus leading-edge genes (defined by requiring |EES| ≥ 3 in at least one of the cohorts) and core pathway information. The seven core thyroid cancer pathways reported earlier here appear depicted in connection with high-consensus leading-edge genes; for simplicity, only genes connected to two or more pathways are shown.

## 4 Discussion

Gene Set Enrichment Analysis (GSEA) pioneered the broad class of functional class scoring approaches to pathway analysis and remains, to this day, one of the leading methods of choice. For this reason, it is of paramount importance for users to be aware of its caveats and limitations. In this Section, we discuss some important aspects of the current implementation of GSEA.

The GSEA algorithm provides a number of alternative statistics that can be used for feature ranking. For categorical phenotypes, options are signal-to-noise (default), t-test, ratio of classes, difference of classes, and log_2_ of ratio of classes. For numerical phenotypes, options are cosine, Euclidean, Manhattan, Pearson and Spearman correlation ranking metrics. These ranking statistics aim to capture some measure of differential expression between categorical phenotypes, or some expression trend associated with quantitative phenotypes. Zyla et al [14] have analyzed these and other ranking metrics, showing that they have a critical impact on GSEA results based on a large number of microarray benchmark datasets. While these metrics are also widely used for RNA-seq datasets, it is important to be aware that these ranking statistics, originally selected for their effectiveness when used with microarray-based expression data, have not been evaluated for their use with data derived from RNA-seq experiments. As noted by Reimand et al [37], rather than using these built-in options, RNA-seq-derived differential gene expression values should be computed outside of GSEA, using methods that include variance stabilization (such as edgeR [48], DESeq [49], limma [33], and voom [50]) and then imported into GSEA software for the actual pathway analysis computation. The current implementation of GSEA, moreover, is not able to handle slightly more complex design matrices, even common ones such as paired data analysis. In this work, we illustrate an approach using paired tumor/non-tumor samples to control for inter-subject heterogeneity, which also makes use of limma’s linear modeling and empirical Bayes moderation to assess differential expression (see Sect. 2.2).

GSEA has two methods for determining the statistical significance of pathway enrichment: gene-set permutation and phenotype permutation (see Sect. 2.4 for details). Gene-set permutation is recommended for studies with limited variability and biological replicates, while phenotype permutation is recommended when a larger number of replicates is available (as a rule of thumb, at least ten per condition) [37]. The main advantage of the latter is that it preserves the structure of gene sets with biologically important gene-gene correlations, in contrast to the gene-set permutation approach [51]. Despite these recommendations, performing phenotype permutation with the appropriate methods that RNA-seq data require for differential expression modeling and ranking is a daunting task for the typical GSEA user, since it demands custom programming to compute pathway enrichment scores and differential expression statistics separately for thousands of phenotype randomizations [37]. Besides the issues of custom programming and computational complexity, phenotype permutation jobs typically take several hours to complete, even in a parallelized multi-CPU environment. By implementing and executing the phenotype permutation approach, however, our work provided evidence that the gene-set permutation method showed superior performance to identify RNA-seq-based, cancer-type-specific pathways.

GSEA offers the option of a weighted Kolmogorov-Smirnov global statistic to assess gene set enrichment; this global statistic is weighed using a power of the local statistic, *p* > 0 (see Sect. 2.4). Besides the so-called classic, or unweighted, original version of GSEA (akin to setting *p* = 0), built-in options for this weight parameter are *p* = 1 (default), *p* = 1.5, and *p* = 2. By increasing *p*, the suppression effects of genes with smaller absolute values of the local statistic become more pronounced, thereby emphasizing the impact of the genes at the top and bottom of the ranked gene list. No clear guidance is offered to users as to how to choose this so-called enrichment statistic with RNA-seq data, since the main evidence in favor of weighted statistics was produced when the method was first introduced for the analysis of microarray datasets [17]. In this work, we explored all available options for both permutation types and concluded that the classic, unweighted gene-set permutation procedure offers comparable or better sensitivity-specificity tradeoffs across cancer types compared with other, more complex and computationally intensive permutation methods. It should be noticed that only performing unweighted GSEA and permuting the genes, analytical solutions for the expected average enrichment score are available [51]; all other GSEA modalities require the calculation of non-parametric, computationally intensive enrichment score distributions.

In fact, by running all GSEA enrichment statistics under gene-set permutation, which can be easily accomplished by uploading a ranked list of genes to GSEA’s Preranked mode, no custom programming is required (except for RNA-seq data processing, QC, normalization, filtering, and differential gene expression analysis, which may be accomplished with standard software outside of GSEA). As demonstrated by our analysis of additional cohorts from non-TCGA sources, we see value in a strategy that uses the proposed Enrichment Evidence Score (EES) as a consensus metric to identify a core set of pathways, complemented by an expanded set of pathways for exploratory analysis, which may be more specifically tailored to the study of interest. Similarly, EES can be used to identify high-consensus leading-edge genes, which allows to integrate pathway- and gene-level information. Hence, the standard, unidirectional sense of information flow through pathway analysis (from genes to pathways) acquires a feedback loop to enrich the input, gene-phenotype associations (e.g. differential gene expression) with functional, gene-pathway associations. The information thus flows from genes to pathways and back.

## 5 Conclusion

The main goal of this work was to quantitatively assess the performance of a variety of GSEA modalities and provide guidance in the practical use of GSEA for the analysis of RNA-seq data. We leveraged harmonized RNA-seq datasets available from TCGA in combination with large, curated pathway collections from MSigDB to obtain cancer-type-specific target pathway lists across multiple cancer types. We performed a detailed analysis of GSEA performance using both gene-set and phenotype permutations combined with four different choices for the Kolmogorov-Smirnov enrichment statistic, either unweighted (also known as classic) or weighted (with three different options for the weight parameter). Based on our benchmarks, we concluded that the classic/unweighted gene-set permutation approach offered comparable or better sensitivity-specificity tradeoffs across cancer types compared with other, more complex and computationally intensive permutation methods, and also appeared superior to the performance of standard over-representation analysis procedures. Finally, we analyzed other large cohorts for thyroid cancer and hepatocellular carcinoma. We utilized a new consensus metric, the Enrichment Evidence Score (EES), which showed a remarkable agreement between pathways identified in TCGA and those from other sources, despite differences in cancer etiology. This procedure suggests an analysis strategy in which, by running multiple GSEA modalities in palallel, one may identify a core set of pathways (namely, those showing a high degree of consensus across enrichment metrics), which may be complemented by an expanded set of pathways for exploratory analysis. The EES metric can also be utilized to identify high-consensus leading-edge genes and integrate pathway- and gene-level information. In accordance to FAIR Principles for Research Software (FAIR4RS) [52], we made all source code openly and publicly available along with detailed step-by-step documentation [27], with the two-pronged aims (i) to ensure the transparency and reproducibility of our results, as well as (ii) to enable further RNA-seq-based benchmarking work by setting up a suitable and flexible framework. We believe that our work fills the existing gap in current guidelines and benchmarks for the use of GSEA with RNA-seq data and that, hopefully, it will stimulate further efforts to advance pathway analysis methods.

## Supporting information

**S1 Table Genes measured and differentially expressed in each TCGA project**. Number of measured protein-coding genes after all filters were applied and how many of them were found differentially expressed in either direction, i.e. over-expressed in primary-tumor (*FC >* 1) or non-tumor (*FC <* 1) tissue, using Bonferroni or Benjamini-Hochberg (B-H) q − value < 0.05 significance thresholds adjusted for multiple testing.

**S2 Table Cancer-type specific pathways**. For each TCGA study, we provide name, size, collection, and GMT file name for all cancer-type specific preselected pathways. Additional columns indicate whether preselected pathways were included in the target and positive-control pathway lists.

**S3 Table Gene ranking scores for all TCGA projects analyzed in our study**.

**S4 Table Enrichment Evidence Scores of thyroid cancer pathways in TCGA-THCA and REBC-THYR**.

**S5 Table Enrichment Evidence Scores of liver cancer pathways in TCGA-LIHC and MO-HCC**.

**S6 Table Enrichment Evidence Scores for genes in the LUI THYROID CANCER CLUSTER 1 pathway in TCGA-THCA and REBC-THYR**.

**S1 Fig. Differentially-expressed genes in TCGA-BRCA**. Volcano plot showing significant genes over-expressed in primary-tumor (red) or non-tumor (blue) tissue based on Bonferroni-adjusted q − value < 0.05. Labels for the top ten genes on either side are also shown.

**S2 Fig. Comparison of gene ranking scores obtained from differential gene expression (DGE) assessed via voom quantile normalization vs log-transformed transcript per million (TPM) normalized counts**. DGE was assessed from 228 paired primary-tumor vs non-tumor breast cancer samples from TCGA-BRCA. The linear fit is shown by a dashed red line. Spearman’s correlation estimate and p-value are shown in the legend.

**S3 Fig. Comparison of pathway ranking scores obtained from differential gene expression (DGE) assessed via voom quantile normalization vs log-transformed transcript per million (TPM) normalized counts**. DGE was assessed from 228 paired primary-tumor vs non-tumor breast cancer samples from TCGA-BRCA. Target pathways were analyzed using GSEA with different enrichment statistics: (a) classic (unweighted); (b) weight parameter *p* = 1; (c) weight parameter *p* = 1.5; (d) weight parameter *p* = 2. Pathway ranking scores were defined as − log_10_(p − value) ∗ sign(ES), where p-values were empirically determined via gene-set permutations and ES represented gene set enrichment scores. Because we used 1000 permutations for GSEA’s null model, we adopted p − value = 0.001 as lower threshold, which implies that pathway ranking scores are constrained to the [ − 3, 3] range. Linear fits are shown by dashed red lines. Spearman’s correlation estimates and p-values are shown in the legends.

**S4 Fig. Jaccard index matrix to assess overlaps among 72 TCGA-BRCA positive control pathways**.

**S5 Fig. Fractional pairwise overlap matrix to assess redundancies among 72 TCGA-BRCA positive control pathways**.

**S6 Fig. Significant TCGA-BRCA positive control pathways across different weight parameter choices**. Gene-set permutation with p − value < 0.001.

**S7 Fig. Comparison of significant TCGA-BRCA target pathways from paired vs unpaired analyses using gene-set permutation approaches for different enrichment statistics**. The significance criterion was p − value < 0.05. (a) Classic (unweighted). (b) Weight parameter *p* = 1. (c) Weight parameter *p* = 1.5. (d) Weight parameter *p* = 2.

**S8 Fig. Comparison of significant TCGA-BRCA target pathways from paired vs unpaired analyses using phenotype permutation approaches for different enrichment statistics**. The significance criterion was p − value < 0.05. (a) Classic (unweighted). (b) Weight parameter *p* = 1. (c) Weight parameter *p* = 1.5. (d) Weight parameter *p* = 2.

**S9 Fig. ROC curves for different GSEA and ORA approaches using 109 TCGA-BRCA target pathways and 1000 randomized negative controls**. (a) Gene-set permutation GSEA. (b) Phenotype permutation GSEA. (c) Signed ORA. (d) Unsigned ORA. GSEA approaches used different enrichment statistics, as indicated. ORA approaches used Bonferroni and Benjamini-Hochberg (B-H) adjusted p-values as different inclusion criteria to select differentially expressed genes, as indicated.

**S10 Fig. AUC across TCGA projects for different GSEA and ORA approaches using cancer-type-specific target pathways and 1000 randomized negative controls**. (a) Gene-set permutation GSEA. (b) Phenotype permutation GSEA. (c) Signed ORA. (d) Unsigned ORA. GSEA approaches used different enrichment statistics, as indicated. ORA approaches used Bonferroni and Benjamini-Hochberg (B-H) adjusted p-values as different inclusion criteria to select differentially expressed genes, as indicated.

**S11 Fig. ROC curves for gene-set permutation GSEA from TCGA-BRCA samples**. Results obtained by an alternative gene expression analysis derived via the edgeR-voom-limma pipeline described in Sect. 2.2.

**S12 Fig. AUC for gene-set permutation GSEA across TCGA projects**. Results obtained by an alternative gene expression analysis derived via the edgeR-voom-limma pipeline described in Sect. 2.2.

### Acknowledgments

This research was supported entirely by the Intramural Research Program of the National Institute on Aging at the U.S. National Institutes of Health.

## References

1. Nguyen TM, Shafi A, Nguyen T, Draghici S. Identifying significantly impacted pathways: a comprehensive review and assessment. Genome Biol. 2019;20(1):1–15.

2. Maleki F, Ovens K, Hogan DJ, Kusalik AJ. Gene set analysis: challenges, opportunities, and future research. Front Genet. 2020;11:654.

3. Xie C, Jauhari S, Mora A. Popularity and performance of bioinformatics software: the case of gene set analysis. BMC Bioinform. 2021;22(1):1–16.

4. Mubeen S, Kodamullil AT, Hofmann-Apitius M, Domingo-Fernández D. On the influence of several factors on pathway enrichment analysis. Brief Bioinform. 2022;23(3):1–13.

5. Liberzon A, Subramanian A, Pinchback R, Thorvaldsdóttir H, Tamayo P, Mesirov JP. Molecular signatures database (MSigDB) 3.0. Bioinformatics. 2011;27(12):1739–40.

6. Khatri P, Sirota M, Butte AJ. Ten years of pathway analysis: current approaches and outstanding challenges. PLoS Comput Biol. 2012;8(2):e1002375.

7. Tarca AL, Bhatti G, Romero R. A comparison of gene set analysis methods in terms of sensitivity, prioritization and specificity. PLoS One. 2013;8(11):e79217.

8. Bayerlová M, Jung K, Kramer F, Klemm F, Bleckmann A, Beißbarth T. Comparative study on gene set and pathway topology-based enrichment methods. BMC Bioinform. 2015;16(1):334.

9. Jaakkola MK, Elo LL. Empirical comparison of structure-based pathway methods. Brief Bioinform. 2016;17(2):336–45.

10. Rahmatallah Y, Emmert-Streib F, Glazko G. Gene set analysis approaches for RNA-seq data: performance evaluation and application guideline. Brief Bioinform. 2016;17(3):393–407.

11. Mathur R, Rotroff D, Ma J, Shojaie A, Motsinger-Reif A. Gene set analysis methods: a systematic comparison. BioData Mining. 2018;11(1):1–19.

12. Ihnatova I, Popovici V, Budinska E. A critical comparison of topology-based pathway analysis methods. PLoS One. 2018;13(1):e0191154.

13. Ma J, Shojaie A, Michailidis G. A comparative study of topology-based pathway enrichment analysis methods. BMC Bioinform. 2019;20(1):1–14.

14. Zyla J, Marczyk M, Domaszewska T, Kaufmann SHE, Polanska J, Weiner J. Gene set enrichment for reproducible science: comparison of CERNO and eight other algorithms. Bioinformatics. 2019;35(24):5146–54.

15. Geistlinger L, Csaba G, Santarelli M, Ramos M, Schiffer L, Turaga N, et al. Toward a gold standard for benchmarking gene set enrichment analysis. Brief Bioinform. 2020;22(1):545–56.

16. Mootha VK, Lindgren CM, Eriksson KF, Subramanian A, Sihag S, Lehar J, et al. PGC-1α-responsive genes involved in oxidative phosphorylation are coordinately downregulated in human diabetes. Nat Genet. 2003;34:267–273.

17. Subramanian A, Tamayo P, Mootha VK, Mukherjee S, Ebert BL, Gillette MA, et al. Gene set enrichment analysis: a knowledge-based approach for interpreting genome-wide expression profiles. Proc Natl Acad Sci. 2005;102(43):15545–50.

18. Timmons JA, Szkop KJ, Gallagher IJ. Multiple sources of bias confound functional enrichment analysis of global -omics data. Genome Biol. 2015;16:186.

19. Wijesooriya K, Jadaan SA, Perera KL, Kaur T, Ziemann M. Urgent need for consistent standards in functional enrichment analysis. PLoS Comput Biol. 2022;18(3):e1009935.

20. The Cancer Genome Atlas Research Network. Comprehensive and integrative genomic characterization of hepatocellular carcinoma. Cell. 2017;169:1327–1341.

21. Kanehisa M, Furumichi M, Sato Y, Ishiguro-Watanabe M, Tanabe M. KEGG: integrating viruses and cellular organisms. Nucleic Acids Res 2021;49(D1):D545–51.

22. Jassal B, Matthews L, Viteri G, Gong C, Lorente P, Fabregat A, et al. The reactome pathway knowledgebase. Nucleic Acids Res. 2020;48(D1):D498–D503.

23. The Gene Ontology Consortium. The Gene Ontology resource: enriching a GOld mine. Nucleic Acids Res 2021;49(D1):D325–34.

24. https://www.gsea-MSigDB.org/gsea/downloads.jsp

25. Thomas GA. The Chernobyl Tissue Bank: integrating research on radiation-induced thyroid cancer. J Radiol Prot. 2012;32(1):N77–80.

26. Candia J, Bayarsaikhan E, Tandon M, Budhu A, Forgues M, Tovuu LO, et al. The genomic landscape of Mongolian hepatocellular carcinoma. Nat Commun. 2020;11(1):4383.

27. https://github.com/juliancandia/GSEARNASeq_Benchmarks

28. https://portal.gdc.cancer.gov

29. Jones W, Greytak S, Odeh H, Guan P, Powers J, Bavarva J, Moore HM. Deleterious effects of formalin-fixation and delays to fixation on RNA and miRNA-Seq profiles. Sci Rep. 2019;9(1):6980.

30. Wehmas LC, Hester SD, Wood CE. Direct formalin fixation induces widespread transcriptomic effects in archival tissue samples. Sci Rep. 2020;10(1):14497.

31. https://www.ncbi.nlm.nih.gov/geo/query/acc.cgi?acc=GSE144269

32. https://docs.gdc.cancer.gov/Data/Bioinformatics_Pipelines/Expression_mRNA_Pip

33. Smyth, GK. Linear models and empirical Bayes methods for assessing differential expression in microarray experiments. Stat Appl Genet Mol Biol. 2004;3:Article3.

34. https://www.gsea-MSigDB.org/gsea/MSigDB

35. Tarca AL, Draghici S, Bhatti G, Romero R. Down-weighting overlapping genes improves gene set analysis. BMC Bioinform. 2012;13:136.

36. Simillion C, Liechti R, Lischer HE, Ioannidis V, Bruggmann R. Avoiding the pitfalls of gene set enrichment analysis with setrank. BMC Bioinform. 2017;18:151.

37. Reimand J, Isserlin R, Voisin V, Kucera M, Tannus-Lopes C, Rostamianfar A, et al. Pathway enrichment analysis and visualization of omics data using g: Profiler, GSEA, Cytoscape and EnrichmentMap. Nat Protoc. 2019;14(2):482–517.

38. Phipson B, Smyth GK. Permutation p-values should never be zero: Calculating exact p-values when permutations are randomly drawn. Stat Appl Gene Mole Biol. 2010;9:39.

39. Manly BFJ. Randomization, Bootstrap and Monte Carlo Methods in Biology, Third Edition. Chapman & Hall, New York. 2007;Ch 6.

40. Tavazoie S, Hughes JD, Campbell MJ, Cho RJ, Church GM. Systematic determination of genetic network architecture. Nat Genet. 1999;22(3):281–5.

41. Huang DW, Sherman BT, Lempicki RA. Bioinformatics enrichment tools: paths toward the comprehensive functional analysis of large gene lists. Nucleic Acids Res. 2009;37(1):1–13.

42. Nikitin A, Egorov S, Daraselia N, Mazo I. Pathway studio—the analysis and navigation of molecular networks. Bioinformatics. 2003;19(16):2155–7.

43. Huang W, Sherman BT, Lempicki RA. Systematic and integrative analysis of large gene lists using DAVID bioinformatics resources. Nat Protoc. 2009;4(1):44–57.

44. Reimand J, Arak T, Adler P, Kolberg L, Reisberg S, Peterson H, et al. g:Profiler-a web server for functional interpretation of gene lists (2016 update). Nucleic Acids Res. 2016;44:W83–89.

45. Krämer A, Green J, Pollard Jr J, Tugendreich S. Causal analysis approaches in Ingenuity Pathway Analysis. Bioinformatics. 2014;30(4):523–30.

46. Pan KH, Lih CJ, Cohen SN. Effects of threshold choice on biological conclusions reached during analysis of gene expression by DNA microarrays. Proc Natl Acad Sci U S A. 2005;102(25):8961–5.

47. Robin X, Turck N, Hainard A, Tiberti N, Lisacek F, Sanchez JC, et al. pROC: an open-source package for R and S+ to analyze and compare ROC curves. BMC Bioinformatics. 2011;12:77.

48. Robinson MD, McCarthy DJ, Smyth GK. edgeR: a Bioconductor package for differential expression analysis of digital gene expression data. Bioinformatics. 2010;26:139–140.

49. Anders S, Huber W. Differential expression analysis for sequence count data. Genome Biol. 2010;11:R106.

50. Law CW, Chen Y, Shi W, Smyth GK. voom: Precision weights unlock linear model analysis tools for RNA-seq read counts. Genome Biol. 2014;15:R29.

51. Tamayo P, Steinhardt G, Liberzon A, Mesirov JP. The limitations of simple gene set enrichment analysis assuming gene independence. Statistical Methods in Medical Research. 2016;25(1):472–487.

52. Barker M, Chue Hong NP, Katz DS, Lamprecht AL, Martinez-Ortiz C, Psomopoulos F, et al. Introducing the FAIR Principles for research software. Sci Data. 2022;9(1):622.

