## Supplementary Figures for "Assessment of Gene Set Enrichment Analysis using curated RNA-seq-based benchmarks"

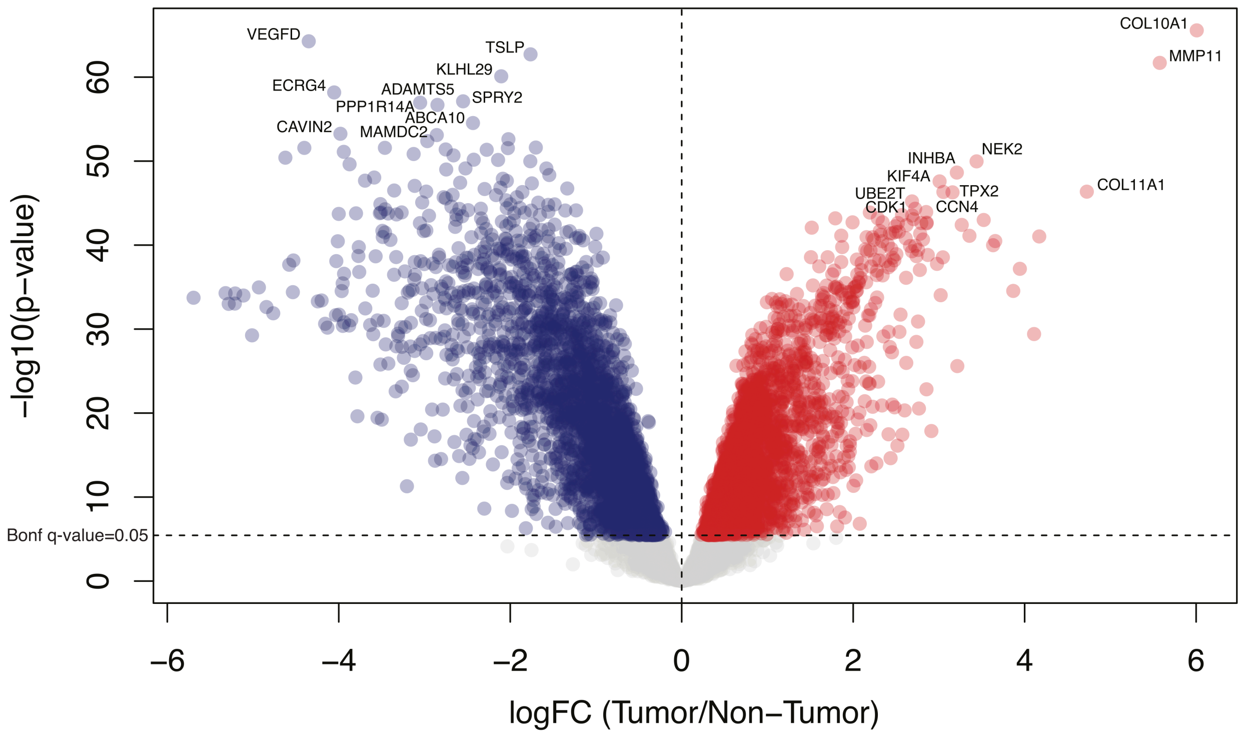


**S1 Fig. Differentially-expressed genes in TCGA-BRCA.** Volcano plot showing significant genes over-expressed in primary-tumor (red) or non-tumor (blue) tissue based on Bonferroni-adjusted q-value<0.05. Labels for the top ten genes on either side are also shown.

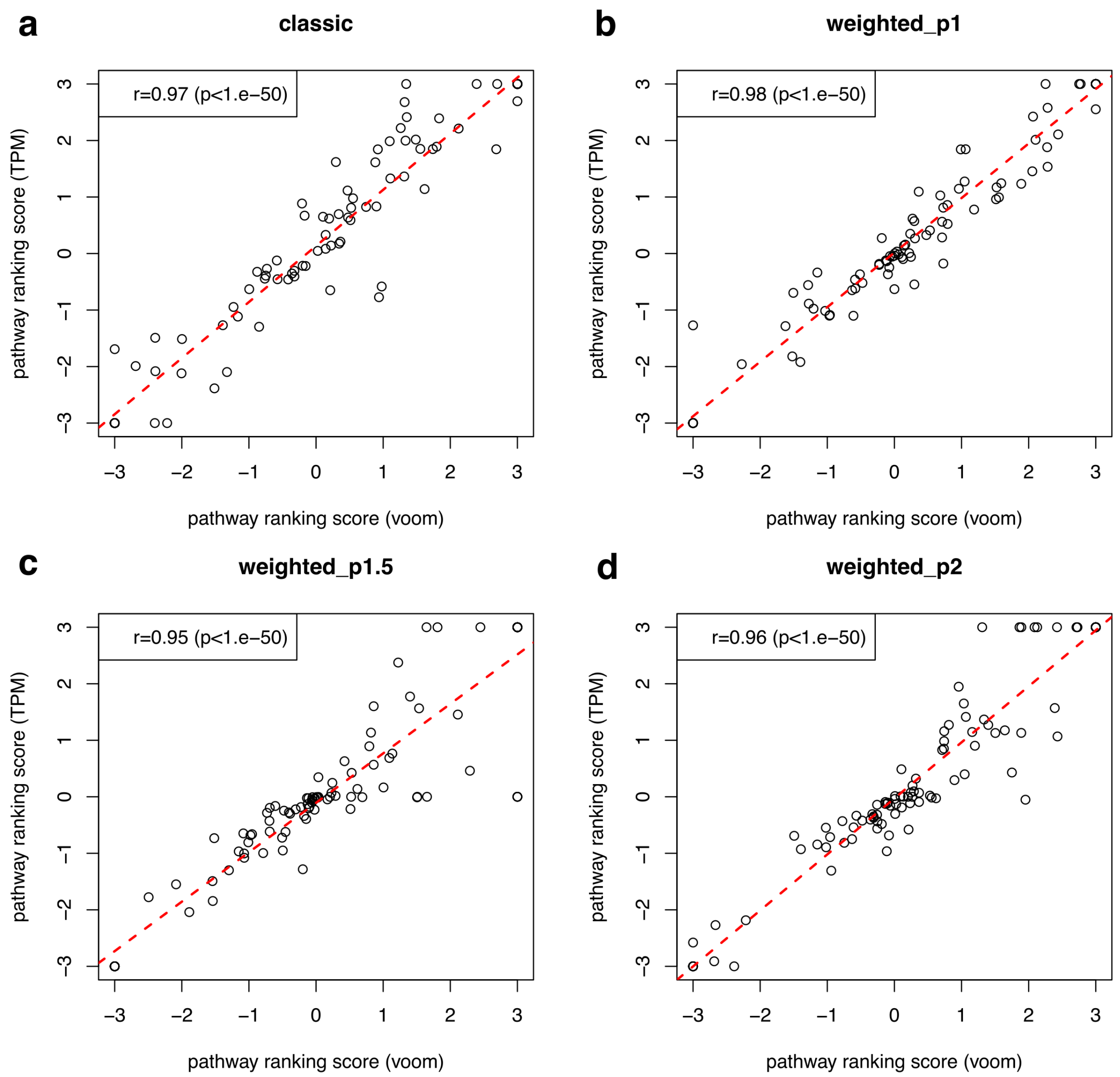


**S3 Fig. Comparison of pathway ranking scores obtained from differential gene expression (DGE) assessed via voom quantile normalization vs log-transformed transcript per million (TPM) normalized counts.** DGE was assessed from 228 paired primary-tumor vs non-tumor breast cancer samples from TCGA-BRCA. Target pathways were analyzed using GSEA with different enrichment statistics: (a) classic (unweighted); (b) weight parameter p=1; (c) weight parameter p=1.5; (d) weight parameter p=2. Pathway ranking scores were defined as -log10(p-value)*sign(ES), where p-values were empirically determined via gene-set permutations and ES represented gene set enrichment scores. Because we used 1000 permutations for GSEA's null model, we adopted p-value=0.001 as lower threshold, which implies that pathway ranking scores are constrained to the [-3,3] range. Linear fits are shown by dashed red lines. Spearman's correlation estimates and p-values are shown in the legends.

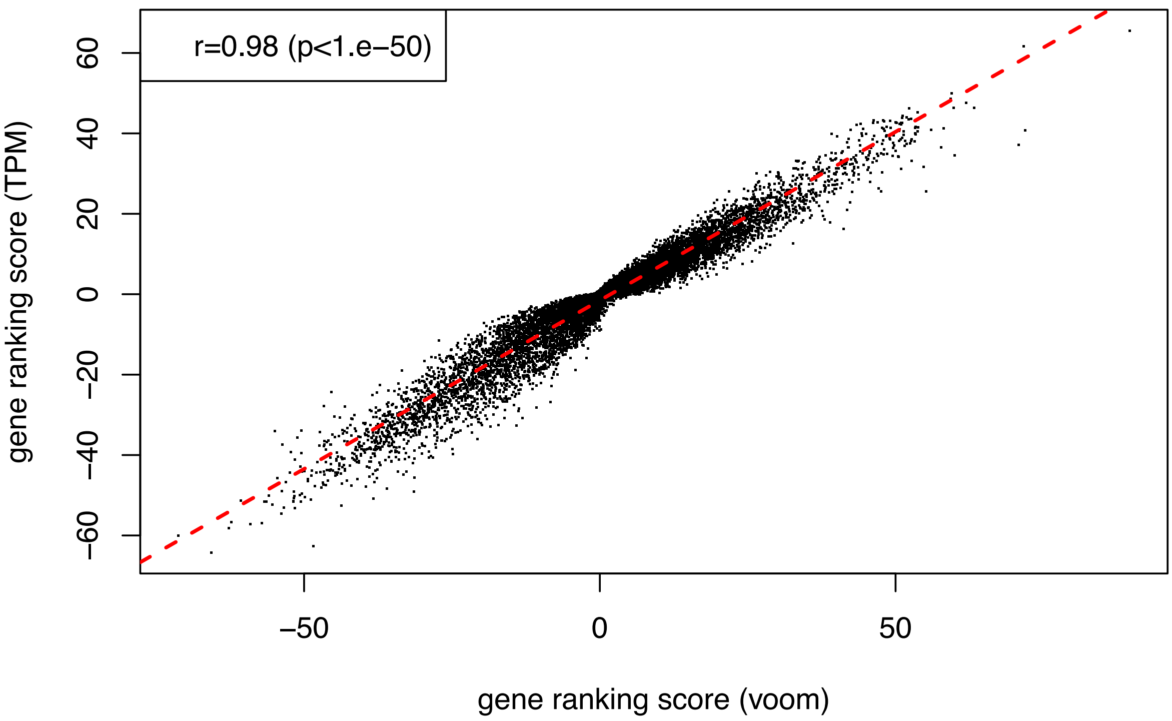


**S2 Fig. Comparison of gene ranking scores obtained from differential gene expression (DGE) assessed via voom quantile normalization vs log-transformed transcript per million (TPM) normalized counts.** DGE was assessed from 228 paired primary-tumor vs non-tumor breast cancer samples from TCGA-BRCA. The linear fit is shown by a dashed red line. Spearman's correlation estimate and p-value are shown in the legend.

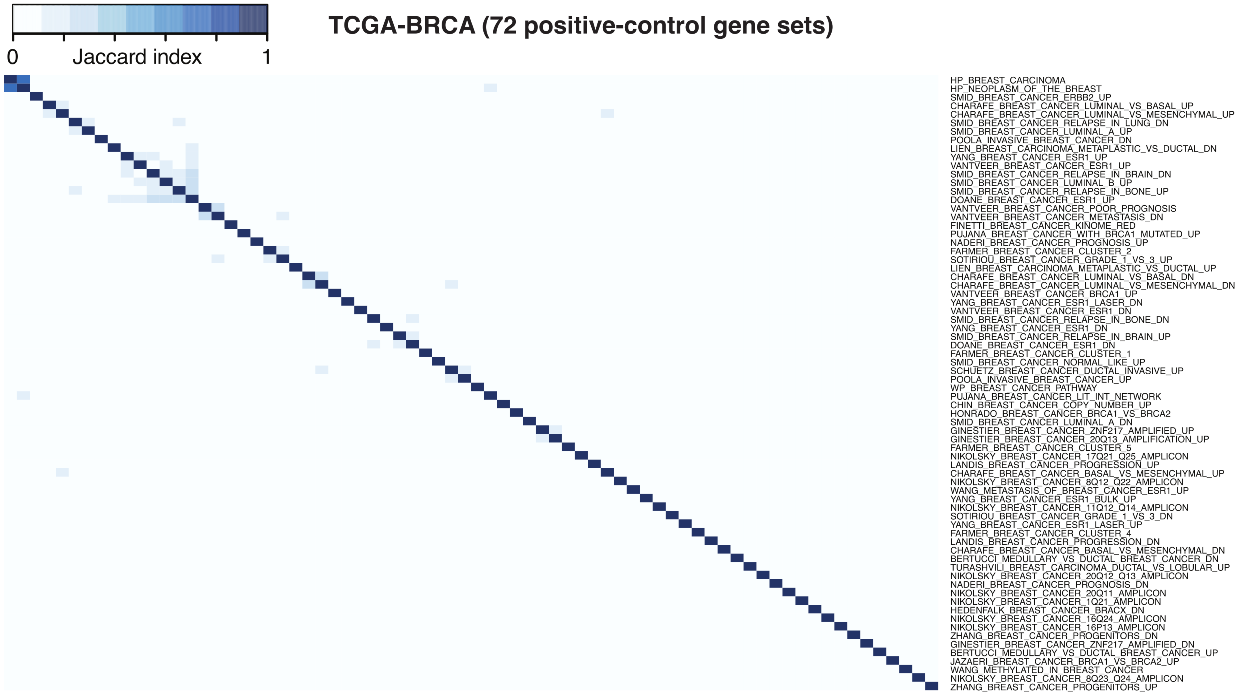


**S4 Fig. Jaccard index matrix to assess overlaps among 72 TCGA-BRCA positive control pathways.**

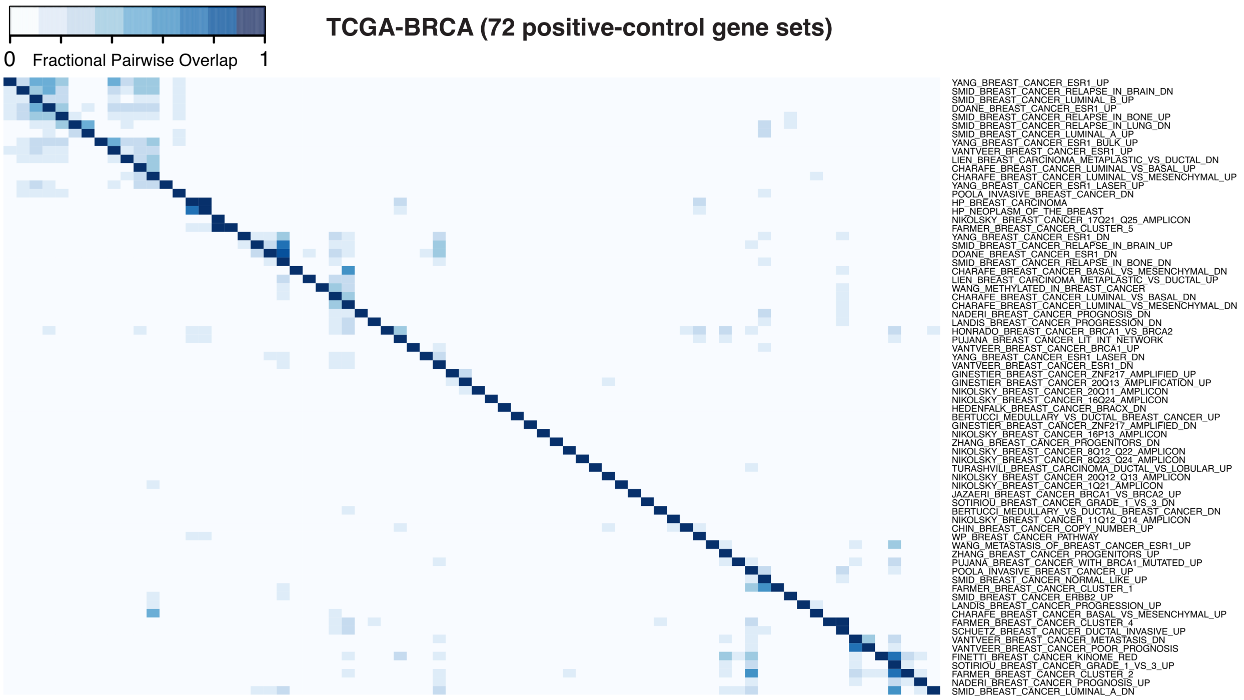


**S5 Fig. Fractional pairwise overlap matrix to assess redundancies among 72 TCGA-BRCA positive control pathways.**

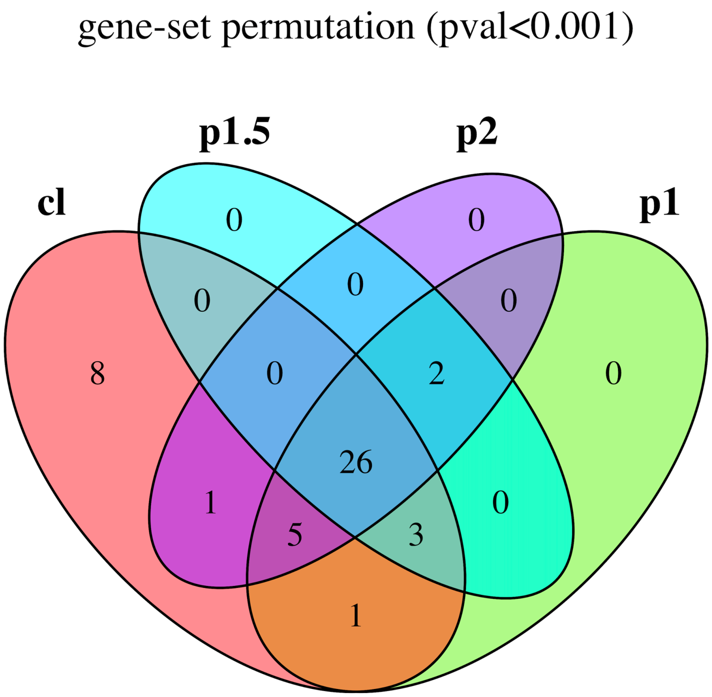


**S6 Fig. Significant TCGA-BRCA positive control pathways across different weight parameter choices.** Gene-set permutation with p-value<0.001.

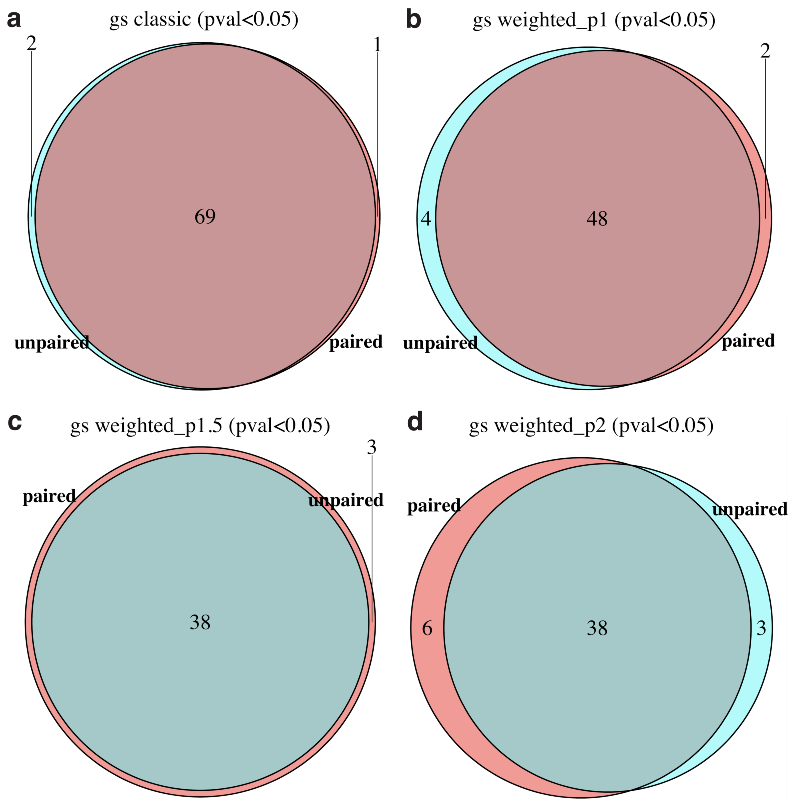


**S7 Fig. Comparison of significant TCGA-BRCA target pathways from paired vs unpaired analyses using gene-set permutation approaches for different enrichment statistics.** The significance criterion was p-value<0.05. (a) Classic (unweighted). (b) Weight parameter p=1. (c) Weight parameter p=1.5. (d) Weight parameter p=2.

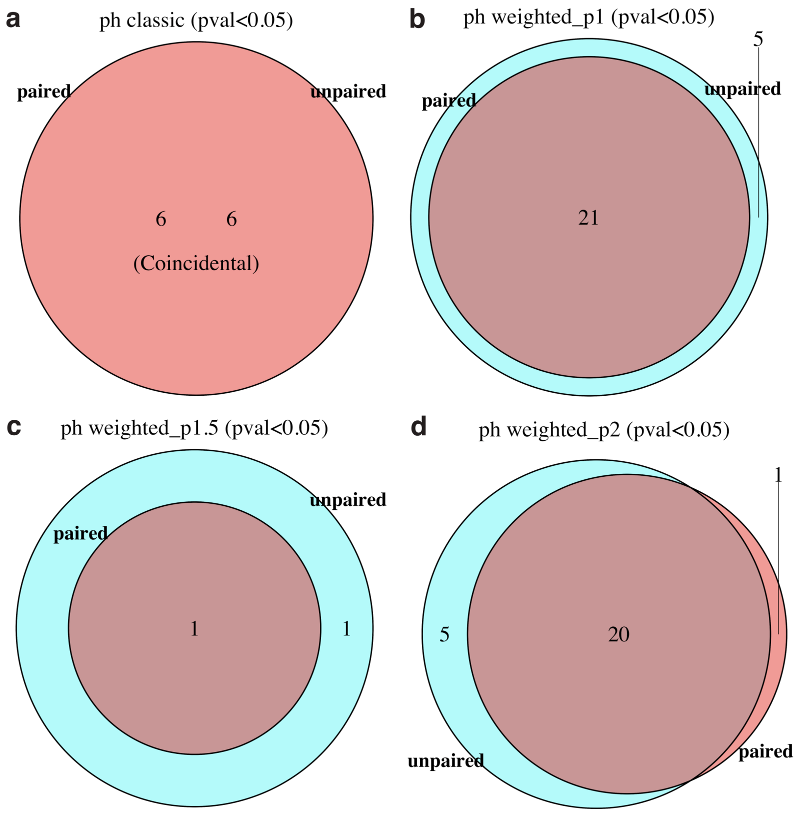


**S8 Fig. Comparison of significant TCGA-BRCA target pathways from paired vs unpaired analyses using phenotype permutation approaches for different enrichment statistics.** The significance criterion was p-value<0.05. (a) Classic (unweighted). (b) Weight parameter p=1. (c) Weight parameter p=1.5. (d) Weight parameter p=2.

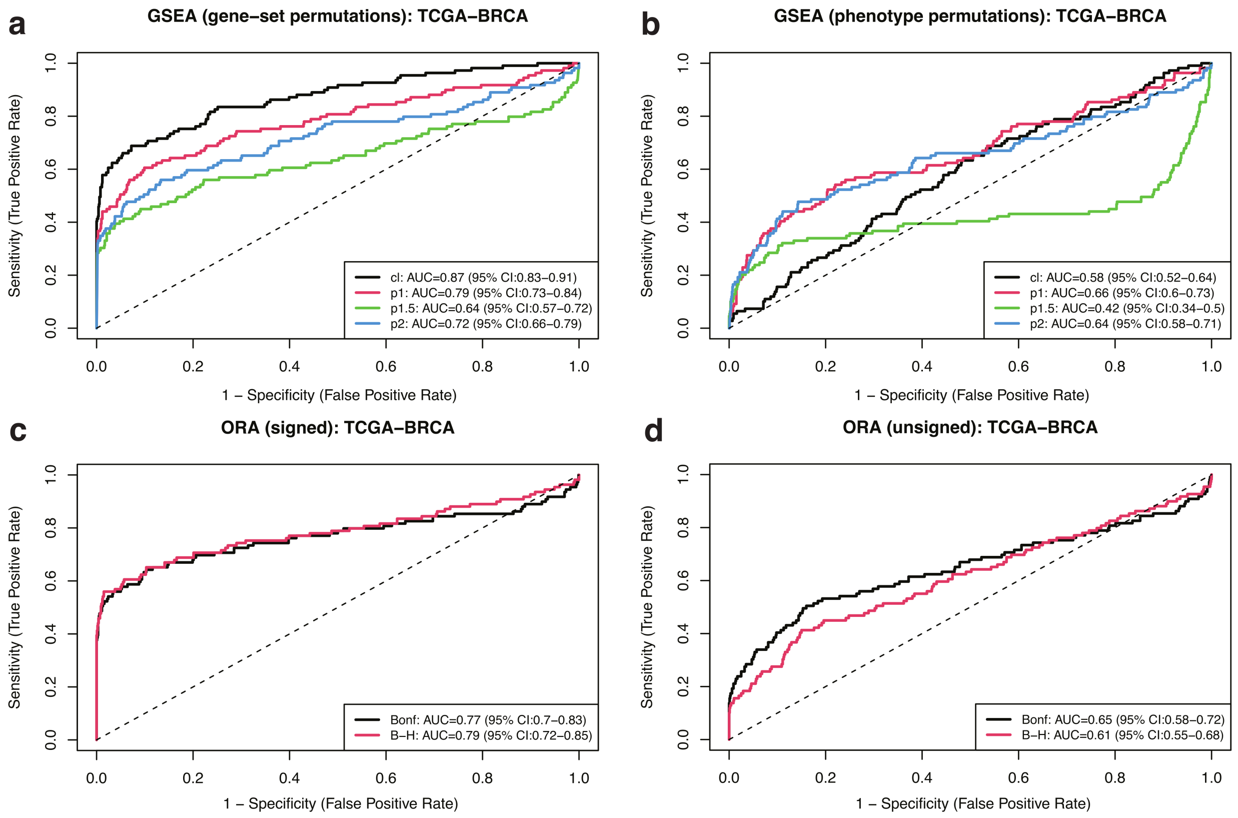


**S9 Fig. ROC curves for different GSEA and ORA approaches using 109 TCGA-BRCA target pathways and 1000 randomized negative controls.** (a) Gene-set permutation GSEA. (b) Phenotype permutation GSEA. (c) Signed ORA. (d) Unsigned ORA. GSEA approaches used different enrichment statistics, as indicated. ORA approaches used Bonferroni and Benjamini-Hochberg (B-H) adjusted p-values as different inclusion criteria to select differentially expressed genes, as indicated.

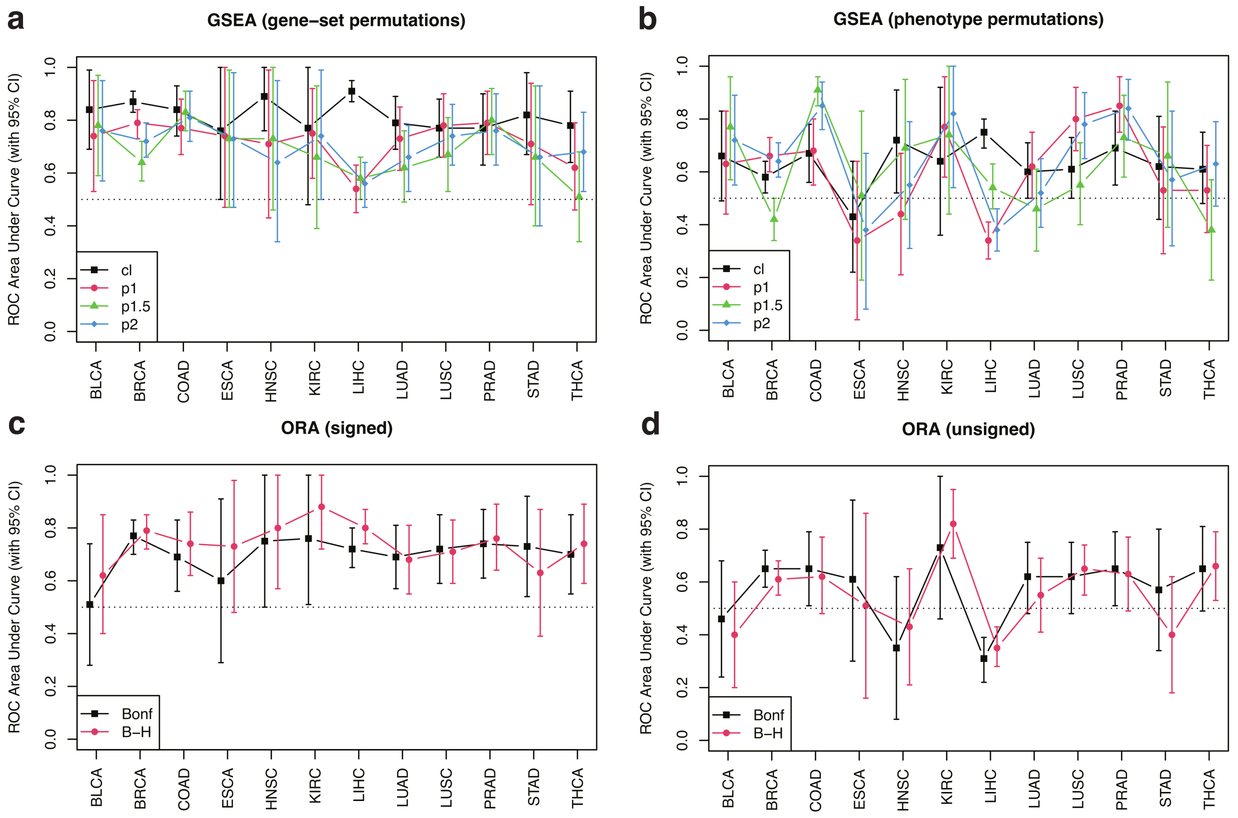


**S10 Fig. AUC across TCGA projects for different GSEA and ORA approaches using cancer-type-specific target pathways and 1000 randomized negative controls.** (a) Gene-set permutation GSEA. (b) Phenotype permutation GSEA. (c) Signed ORA. (d) Unsigned ORA. GSEA approaches used different enrichment statistics, as indicated. ORA approaches used Bonferroni and Benjamini-Hochberg (B-H) adjusted p-values as different inclusion criteria to select differentially expressed genes, as indicated.

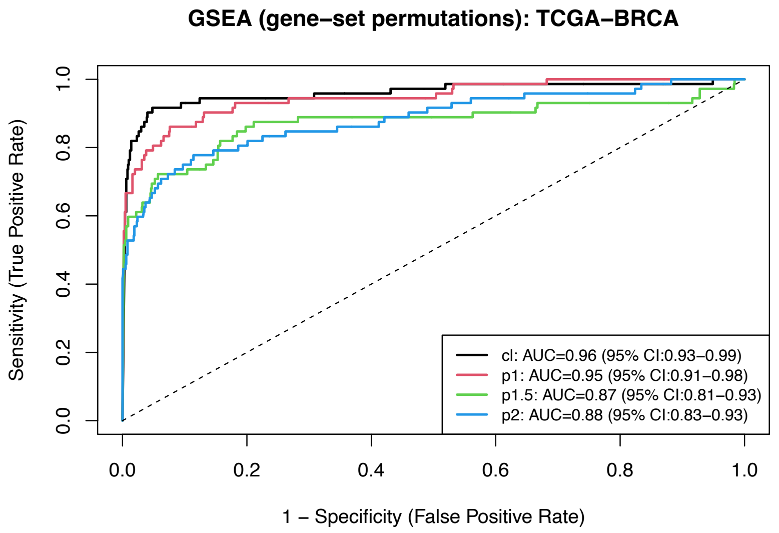


**S11 Fig. ROC curves for gene-set permutation GSEA from TCGA-BRCA samples.** Results obtained by an alternative gene expression analysis derived via the edgeR-voom-limma pipeline described in Sect. 2.2.

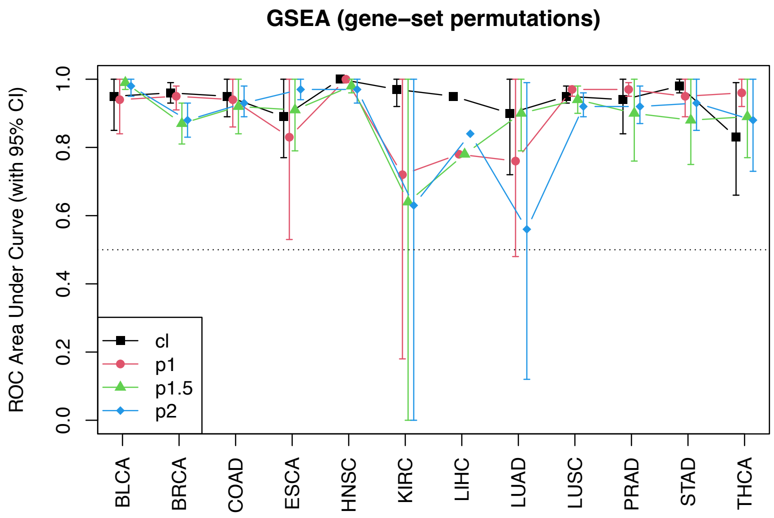


**S12 Fig. AUC for gene-set permutation GSEA across TCGA projects.** Results obtained by an alternative gene expression analysis derived via the edgeR-voom-limma pipeline described in Sect. 2.2.
